# LANCE-seq: Live-tissue Analysis via Needle-encoded Capture Extraction for Spatial Transcriptomics

**DOI:** 10.64898/2026.11.14.766625

**Authors:** Wenjia Wang, Handong Wang, Liyong He, Kaitong Dang, Liuxi Chu, Chen Gao, Kaiqiang Ye, Xiangwei Zhao

**Author notes:** These authors contributed equally (Wenjia Wang, Handong Wang).

## Abstract

Spatial transcriptomics enables the resolution of spatial heterogeneity in gene expression within tissues, yet most existing sequencing-based methods rely on ex vivo tissue processing, making it difficult to track dynamic molecular changes in the same living individual. We developed LANCE-seq (Live-tissue Analysis via Needle-encoded Capture Extraction sequencing), a live-tissue spatial transcriptomics platform based on encoded microneedle arrays that enables high-density spatial RNA capture through poly(amidoamine) (PAMAM) dendrimer-functionalized interfaces and microfluidic orthogonal combinatorial encoding. In frozen mouse brain tissue, LANCE-seq detected an average of 6,399 genes per spatial spot and reconstructed the spatial distribution of major brain regions and representative marker genes. Furthermore, in the acetaminophen (APAP)-induced acute liver injury model, LANCE-seq enabled repeated sampling from the same living animal at different time points. At 6 h after APAP treatment, injury-related genes such as *Hmox1*, *Ddit3*, *Atf3*, and *Fgf21* were significantly elevated, and transcriptional programs associated with stress, inflammation, and cellular injury were broadly enhanced. Meanwhile, liver zonation-related transcriptional signatures were remodeled, with different relative zonation states all exhibiting activation of injury programs. By integrating high-throughput spatial encoding with minimally invasive in vivo sampling, LANCE-seq provides a platform for longitudinal spatial transcriptomic studies of drug responses and disease progression.

---

Spatially resolved transcriptomics (ST) enables high-throughput profiling of gene expression while preserving spatial information, providing a powerful approach for resolving tissue architecture, cellular composition, and microenvironmental heterogeneity^1–3^. Currently, next-generation sequencing (NGS)-based ST technologies^4–6^, represented by spatial barcode array sequencing, have been widely applied in tissue atlas construction and disease research. However, most existing mainstream ST methods largely rely on ex vivo sample processing workflows involving tissue sectioning, fixation, permeabilization, or freezing. Although this *ex vivo*-dependent preparation approach has achieved significant advances in spatial resolution^7–9^, it inevitably strips away the tissue’s native physiological state and complex three-dimensional microenvironment, capturing only a "static snapshot" of the tissue at the moment of fixation^10–12^. Biological processes exhibit continuous temporal evolution. However, in dynamic contexts such as drug response, acute tissue injury, disease progression, and tissue repair, existing ST studies typically require sampling from different animals or different tissue specimens at distinct time points, making it difficult to directly resolve longitudinal spatial transcriptional changes within the same living individual. Therefore, achieving repeated spatial transcriptomic sampling of living tissues while minimizing tissue damage represents a critical technical need for further expanding the temporal dimension of spatial omics^13^.

Microneedle arrays (MNAs) can penetrate tissue barriers with limited tissue damage and have been used for in situ biomarker extraction, biofluid sampling, and multi-omics analysis, highlighting their potential as molecular sampling interfaces for living tissues^14–16^. However, existing needle-based molecular sampling strategies still struggle to simultaneously achieve transcriptome coverage and spatial sampling density. Some studies perform homogenized bulk analysis on microneedle-collected materials and therefore fail to preserve spatial heterogeneity within tissues, whereas other approaches are limited to a small set of preset targets and cannot provide whole-transcriptome measurements^17, 18^. Recently, MSN-seq demonstrated the feasibility of needle-based tissue extraction for spatial transcriptomic measurements by using a needle to perform targeted sampling of specific regions in frozen tissue sections, combined with improved Smart-seq2 for full-length transcriptome analysis^19^. However, this method still relies on ex vivo tissue sections and point-by-point sampling, making it difficult to achieve continuous monitoring of the same living individual. Another study, aivST^12^, further utilized DNA barcode-modified needle arrays to perform repeated transcriptomic sampling in living mice, demonstrating the potential of needle-based devices for longitudinal spatial transcriptomic research. However, the number of spatial sampling units remains limited, making high-density spatial coverage and resolution across a two-dimensional tissue area difficult to achieve. Ji et al. developed Spectrum-FISH, combining nanoneedle molecular fishing with fluorescence-encoded detection to profile mRNAs and post-transcriptional targets in fresh tissue slices and human biopsies at single-cell or subcellular resolution^20^. Although sequencing-free, cost-efficient and spatially high-resolution, it relies on predefined target panels, and its temporal analyses compare excised tissues across developmental stages rather than repeatedly sampling the same living animal, limiting unbiased whole-transcriptome and longitudinal in vivo applications.

Taken together, these studies establish complementary advances in needle-based spatial molecular profiling, but integrating repeated in vivo sampling with whole-transcriptome coverage and high-density two-dimensional spatial encoding remains challenging. Achieving this combination requires improvements in both molecular capture efficiency and scalable spatial encoding. Compared with conventional two-dimensional slides, microneedle arrays possess complex three-dimensional topological structures, where the curved surfaces of the needle bodies and the needle tips increase the difficulty of surface functionalization, uniform probe loading, and microfluidic alignment and sealing^21–23^. On the other hand, the limited tissue volume that can be acquired by a single microneedle makes the efficiency of RNA capture, reverse transcription, and library construction under low-input conditions a key constraint^24, 25^. When spatial sampling expands from a small number of needles to thousands of sites, it is also necessary to assign resolvable two-dimensional coordinates to each sampling unit and control crosstalk between adjacent coding regions^26, 27^. Therefore, simultaneously achieving high-density probe loading, high-fidelity spatial encoding, and stable molecular capture on a three-dimensional microneedle surface is a core challenge in building a high-throughput live-tissue spatial transcriptomics platform.

Based on these requirements, we developed LANCE-seq (Live-tissue Analysis via Needle-encoded Capture Extraction sequencing), a live-tissue spatial transcriptomics platform based on encoded microneedle arrays. By constructing a poly(amidoamine) (PAMAM) dendrimer functional interface on the surface of silicon-based microneedles, combined with a microfluidics-assisted orthogonal combinatorial encoding strategy, we built 2,500 RNA capture units with independent two-dimensional spatial coordinates on a 50 × 50 microneedle array. First, we systematically evaluated the transcriptome capture performance, technical reproducibility, and spatial structure reconstruction capability of LANCE-seq using frozen mouse brain tissue. Subsequently, we applied this platform to an acetaminophen (APAP)-induced acute liver injury model, analyzing changes in transcriptional states and their spatial distributions before and after drug treatment through repeated microneedle sampling at different time points from the same living animal. LANCE-seq, by integrating high-density spatial encoding with minimally invasive tissue sampling, offers a novel technical strategy for longitudinal spatial transcriptomic analysis in living tissues.

## Results

### Design and Workflow of LANCE-seq

To achieve *in situ* capture of the spatial transcriptome on three-dimensional substrates with tissue-penetrating capability, we constructed the LANCE-seq platform. The core device is positioned in the central region of a standard 75 × 25 mm silicon substrate and consists of a 50 × 50 array of silicon microneedles, providing 2,500 sampling units in total. Each individual microneedle has a base diameter of 50 μm, a height of 100 μm, and a center-to-center spacing of 200 μm. Fig. 1a illustrates the overall design of LANCE-seq and the spatial transcriptome library construction workflow. For the functionalization of the microneedle surface, we adopted the surface chemistry strategy from Decoder-seq, constructing a three-dimensional PAMAM dendrimer functional layer on the silicon-based microneedle surface (Fig. 1b). Compared with the conventional 3-aminopropyltriethoxysilane (APTES)-modified two-dimensional surface, the PAMAM-modified surface exhibited stronger fluorescence signals from complementary probe hybridization, suggesting that the three-dimensional dendritic structure can enhance the loading capacity of surface DNA probes. After alkali treatment, the fluorescence signals essentially disappeared, with no obvious nonspecific fluorescence residue observed (Fig. 1c).

**Fig. 1.**
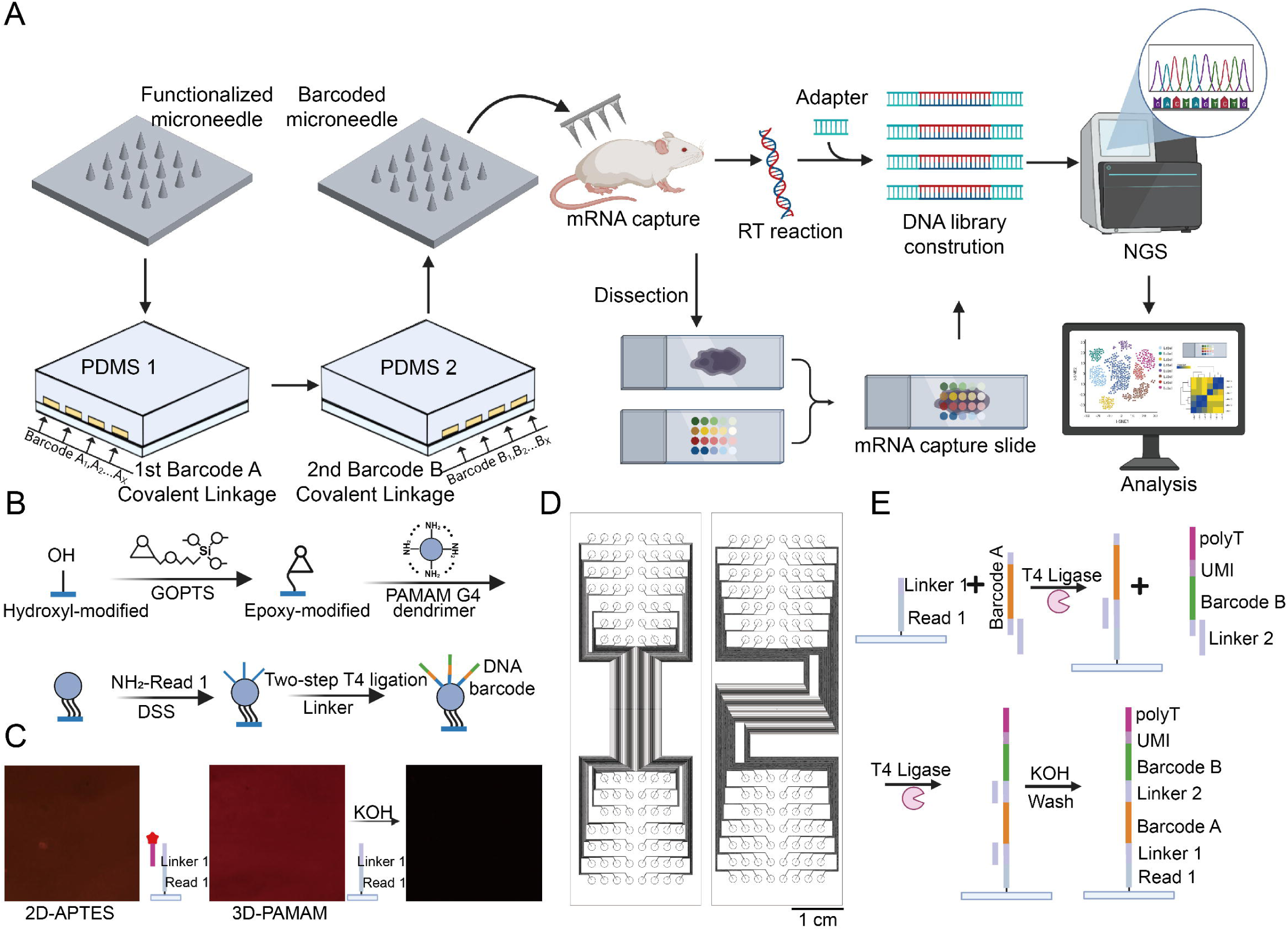
Design and workflow of the LANCE-seq platform. (a) Schematic overview of the LANCE-seq platform, showing minimally invasive microneedle penetration for *in situ* RNA capture and the workflow for spatial transcriptomic library construction. (b) Schematic of surface functionalization showing formation of a 3D PAMAM dendrimer layer on the silicon microneedle array. (c) Fluorescence characterization comparing complementary-probe hybridization signals on 3D dendrimer-modified microneedles and conventional 2D APTES-modified glass slides. The dendrimer-modified microneedles showed higher fluorescence intensity, and the signal was largely removed after alkaline washing. (d) Design of the customized PDMS microfluidic channels (100 μm in both width and spacing) utilized for spatial encoding. (e) Illustration of the orthogonal combinatorial encoding strategy, showing sequential *in situ* T4 ligation of Barcode A and Barcode B probes to generate 2,500 capture spots with unique physical coordinates (X, Y) on the array.

To achieve high-throughput two-dimensional spatial encoding of the microneedle array, we employed an orthogonal combinatorial encoding strategy assisted by a custom-designed polydimethylsiloxane (PDMS) microfluidic chip. Both the microfluidic channel width and the channel spacing were 100 μm (Fig. 1d). After pre-modifying the substrate with Read 1 probes via covalent conjugation, two sets of mutually perpendicular PDMS microfluidic channels were sequentially aligned with the microneedle array, and 50 types of Barcode A and 50 types of Barcode B were introduced for two rounds of T4 ligation reactions, respectively. The combination of the two barcode sets ultimately generated 2,500 capture units with defined two-dimensional coordinates (Fig. 1e). Following the spatial encoding, the microneedle array was pressed into the target tissue for in situ RNA capture. After array retrieval, tissue permeabilization, reverse transcription, and second-strand synthesis were sequentially performed on the microneedle surface. The cDNA carrying spatial barcodes was then released via alkaline elution, followed by PCR amplification, sequencing adapter ligation, and high-throughput sequencing to generate the spatial transcriptomic library (Fig. 1a).

### Optimization of Orthogonal Spatial Encoding

To achieve optimal probe loading efficiency and validate the reliability of the spatial encoding strategy, we systematically optimized and characterized the biochemical reaction conditions for platform construction. First, to determine the optimal concentration of Read 1 probes crosslinked to the substrate, we evaluated substrates modified with 5–50 μM Read 1 probes. Hybridization with complementary fluorescent probes and quantitative analysis revealed that the fluorescence signal in the modified regions gradually increased with increasing probe concentration and approached saturation at 40 μM (Fig. 2a). Subsequently, we precisely aligned and pressed PDMS microchannels against the rows and columns of the microneedle array under a microscope and sequentially performed the directional ligation of Barcode A and Barcode B (Fig. 2b). Optimization of the Barcode A ligation concentration (2–30 μM) revealed a highly uniform fluorescence signal distribution across the microchannels, with no obvious fluorescence leakage or cross-flow between adjacent channels. Considering both modification efficiency and probe cost, 10 μM was determined to be the optimal Barcode A ligation concentration (Fig. 2c).

**Fig. 2.**
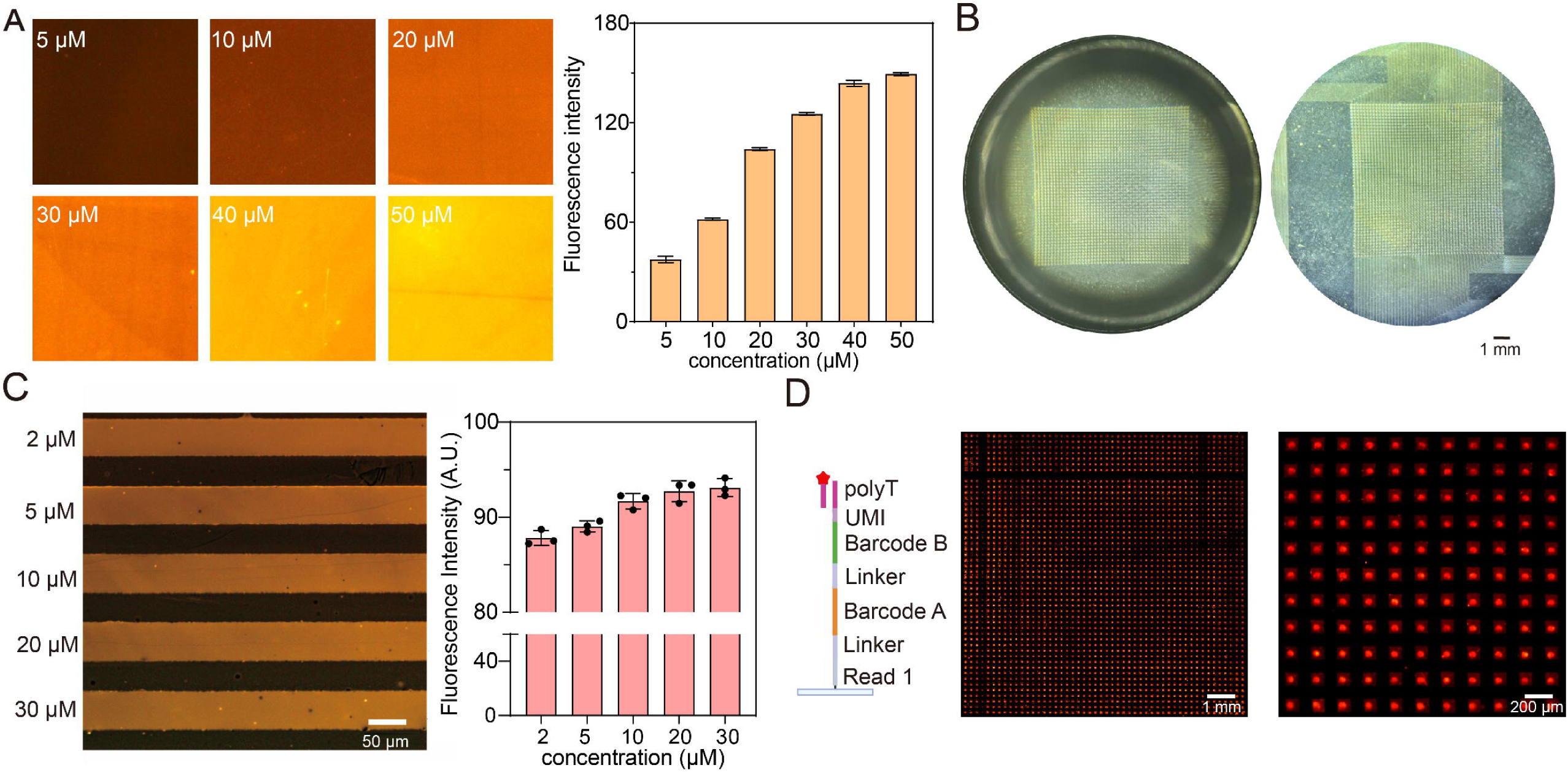
Optimization and validation of orthogonal spatial encoding. (a) Quantification of fluorescence intensity across substrate-conjugated Read 1 probe concentrations of 5–50 μM. The fluorescence intensity increased and reached saturation at 40 μM. (b) Microscopic images demonstrating the precise alignment and firm clamping of the PDMS microfluidic channels with the rows and columns of the microneedle array for directed probe ligation. (c) Optimization of the Barcode A ligation concentration (2–30 μM). The microchannels showed highly uniform fluorescence with no observable leakage or crosstalk. A concentration of 10 μM was selected for subsequent experiments. (d) Global fluorescence characterization of the 50 × 50 combinatorially encoded array (2,500 spatial spots). The fluorescence pattern showed distinct square boundaries, with stronger signals concentrated on the microneedle bodies (central circular regions). Data are meanß±ßs.e.m.

Based on the above optimization conditions, we further completed the spatial combinatorial encoding of the microneedle array. After hybridization with complementary fluorescent probes, the array exhibited a regular 50 × 50 spatial encoding pattern, generating a total of 2,500 independent spatial sites (Fig. 2d). The boundaries of each encoding region were clearly defined, with no obvious cross-talk observed between adjacent regions, and the strong fluorescent signals were primarily concentrated in the microneedle body regions at the center of each spatial unit. To further evaluate the modification uniformity of the spatially encoded probes on the three-dimensional microneedle surface, we performed fluorescence imaging at the focal planes of both the microneedle tips (Top) and the substrate (Bottom) of the array (Supplementary Fig. 1a). All regions of the array exhibited stable fluorescence signal distribution (Supplementary Fig. 1b), indicating that the combinatorial encoding strategy enables relatively uniform probe modification across the entire microneedle array. Meanwhile, the fluorescence intensity in the microneedle body regions was overall higher than that in the relatively flat substrate regions, suggesting that the needle body regions can achieve stronger probe-associated fluorescence signals. These results validate the feasibility of orthogonal microfluidic combinatorial encoding on microneedle arrays, laying the foundation for subsequent spatial capture of tissue RNA.

### LANCE-seq Recapitulates Spatial Transcriptomic Architecture in Frozen Mouse Brain

To evaluate the transcriptome capture performance of LANCE-seq in tissue samples, we utilized microneedle arrays for in situ sampling and validation on frozen ex vivo mouse brain tissue (Fig. 3a). Using conventional two-dimensional planar slide capture (Section group) as a benchmark, we compared the transcriptome complexity obtained from the microneedle platform (MN group). Violin plots revealed that at a sequencing saturation of 46.9%, the Section group detected an average of 4,837 genes and 32,051 unique molecular identifiers (UMIs) per spatial spot, whereas the MN group, at a lower sequencing saturation of 36.4%, still detected an average of 6,399 genes and 25,063 UMIs per spatial spot (Fig. 3b). These results indicate that microneedle sampling yields high-complexity transcriptomic profiles suitable for downstream spatial transcriptomic analysis. In addition, the proportion of reads with valid spatial barcodes in the MN group was lower than that in the Section group (Fig. 3c). Global gene expression correlation analysis revealed that the MN group and the Section group maintained a high correlation at the overall expression level, with a Pearson correlation coefficient of 0.872 (Fig. 3d). The correlation between two technical replicates derived from the same brain tissue processed by independent microneedle arrays was even higher, with a Pearson correlation coefficient reaching 0.961 (Fig. 3e), indicating that LANCE-seq exhibits good technical reproducibility.

**Fig. 3.**
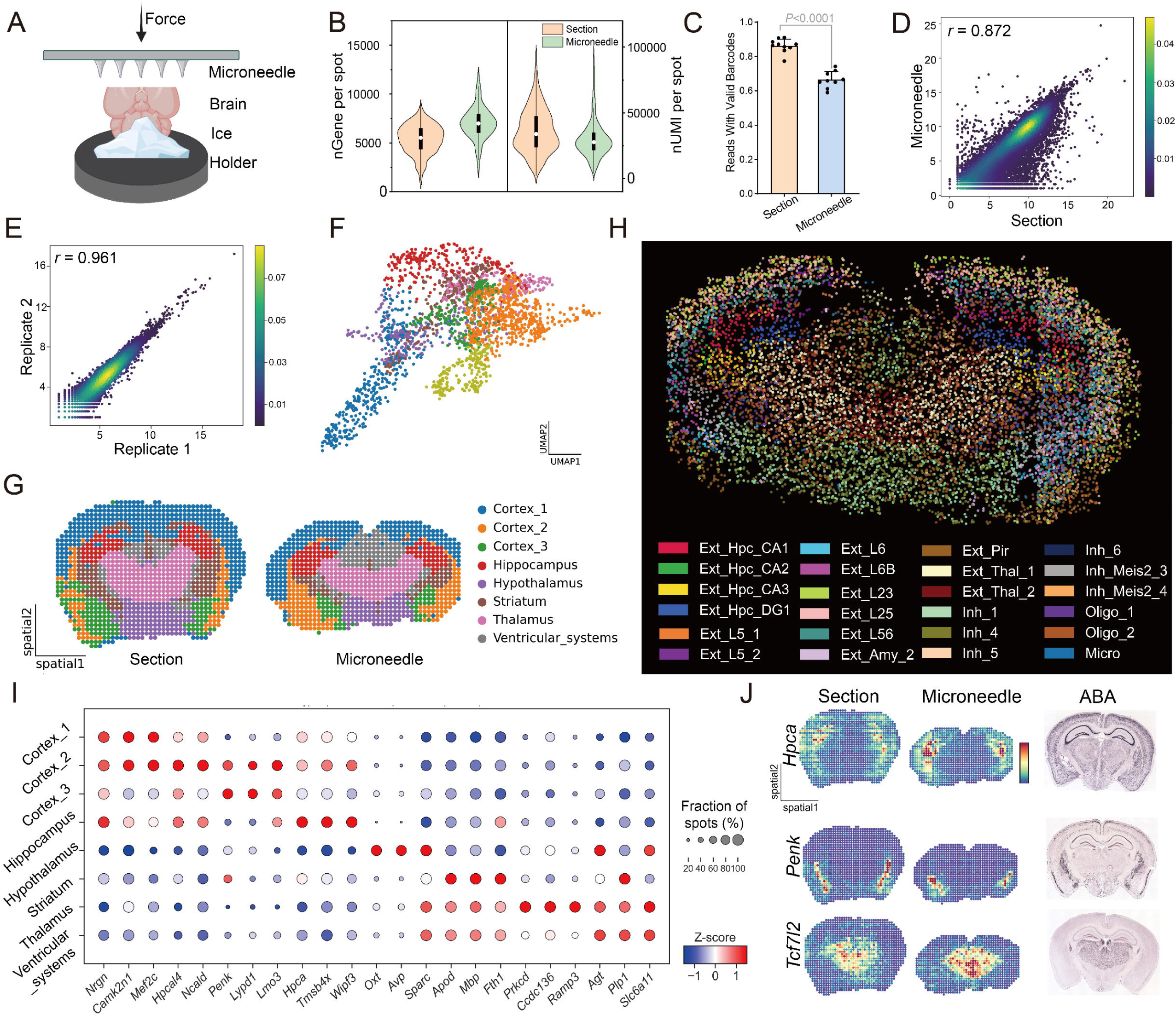
Validation of LANCE-seq in frozen mouse brain tissue. (a) Schematic illustration of *in situ* sampling on frozen *ex vivo* mouse brain tissues using the microneedle arrays. (b) Violin plots comparing the numbers of detected genes and UMI counts per spatial spot between the Microneedle (MN) and conventional planar Section groups. (c) Comparison of the fraction of reads with valid barcodes between the MN (*n* = 9) and Section (*n* = 10) groups, error bar indicates meanß±ßs.d. (d) Global gene expression correlation analysis between the MN and Section groups (*r* = 0.872). (e) Technical reproducibility analysis between independent microneedle-array measurements from the same brain tissue source (*r* = 0.961). (f) UMAP visualization of the spatial transcriptomic data, colored by the eight transcriptional clusters identified by unsupervised clustering. (g) Spatial mapping of the identified clusters, reconstructing major anatomical regions of the mouse brain. (h) Spatial cell-type deconvolution maps showing the inferred localization and enrichment patterns of 24 representative neuronal and glial subpopulations on the microneedle array. (i) Bubble plot showing representative marker genes across the anatomically annotated regions. The top three marker genes for each anatomically annotated region were selected for visualization. Bubble color represents the gene-wise Z-score of mean expression across anatomical regions, whereas bubble size represents the fraction of spatial spots expressing the corresponding gene within each region. (j) Spatial expression patterns of representative marker genes (*Hpca*, *Penk* and *Tcf7l2*) in the MN and Section datasets and in Allen Brain Atlas (ABA) in situ hybridization (ISH) reference data.

Furthermore, we reconstructed the spatial transcriptomic atlas of the mouse brain. Through joint dimensionality reduction and unsupervised clustering analysis (Fig. 3f, Supplementary Fig. 2), we identified eight major transcriptional clusters in the merged data. Mapping these clusters back to the tissue space, both groups recapitulated spatial distributions corresponding to the anatomical locations of major brain regions, including the cortex (Cortex 1–3), hippocampus (Hippocampus), hypothalamus (Hypothalamus), striatum (Striatum), thalamus (Thalamus), and ventricular systems (Ventricular systems) (Fig. 3g). Using mouse whole-brain single-nucleus RNA sequencing data as a reference, we deconvolved the cellular composition at each spatial spot of the microneedle array and inferred the relative abundances and spatial enrichment patterns of 24 representative neuronal and glial cell subtypes. Excitatory neurons such as Ext L23/L5/L6, inhibitory neurons such as Inh_1/Inh_4, and glial cell subtypes such as Oligo and Micro exhibited spatial distributions generally consistent with known brain tissue anatomical structures (Fig. 3h).

The marker gene expression bubble plot further supported the above anatomical region annotations, with different regions exhibiting enriched expression of their corresponding representative marker genes, such a*s Mef2c* and *Camk2n1* in the cortex, *Hpca* in the hippocampus, *Oxt* and *Avp* in the hypothalamus, *Apod* and *Mbp* in the striatum, and *Agt* and *Plp1* in the ventricular system (Fig. 3i). Finally, we compared the spatial expression patterns of representative brain region marker genes, including *Hpca*, *Penk*, and *Tcf7l2*. The regional expression patterns of these genes in the MN group were generally consistent with those observed in the Section group and with the in situ hybridization reference data from the Allen Brain Atlas (ABA)^28^ (Fig. 3j). Compared with the Section group, some brain region boundaries reconstructed in the MN group were relatively diffuse, but the major brain regions and their representative transcriptional signatures remained clearly identifiable. In summary, LANCE-seq is capable of obtaining spatially informative transcriptomic data from complex tissues and reconstructing the spatial expression patterns of major tissue regions and marker genes, thereby validating its capacity for spatial transcriptomic profiling in complex tissues.

### LANCE-seq Enables Longitudinal In Vivo Transcriptomic Sampling of the Mouse Liver

We further evaluated its applicability in repeated sampling of living tissues. To this end, we established an APAP-induced acute liver injury model in mice and employed four mice to construct a longitudinal sampling and complementary control design (Fig. 4a). Mouse A underwent microneedle sampling at 0 h before APAP treatment and at 6 h after treatment, yielding MN1 and MN2, respectively, thereby constituting a longitudinal sampling sequence from the same living animal. Mouse B underwent laparotomy and abdominal closure at 0 h without microneedle sampling and received phosphate-buffered saline (PBS). At 6 h, MN3 and CON1 were collected, and Mouse B served as the sham-operated control. Mouse C received APAP at 0 h without prior surgery or microneedle sampling. At 6 h, MN4 and CON2 were collected to evaluate the potential impact of the 0 h surgery and microneedle sampling procedures on the subsequent APAP response. Mouse D had MN5 and CON3 collected under healthy conditions, serving as an independent healthy control. The sample identities, treatments, sampling time points, and experimental roles are summarized in Supplementary Table 3.

**Fig. 4.**
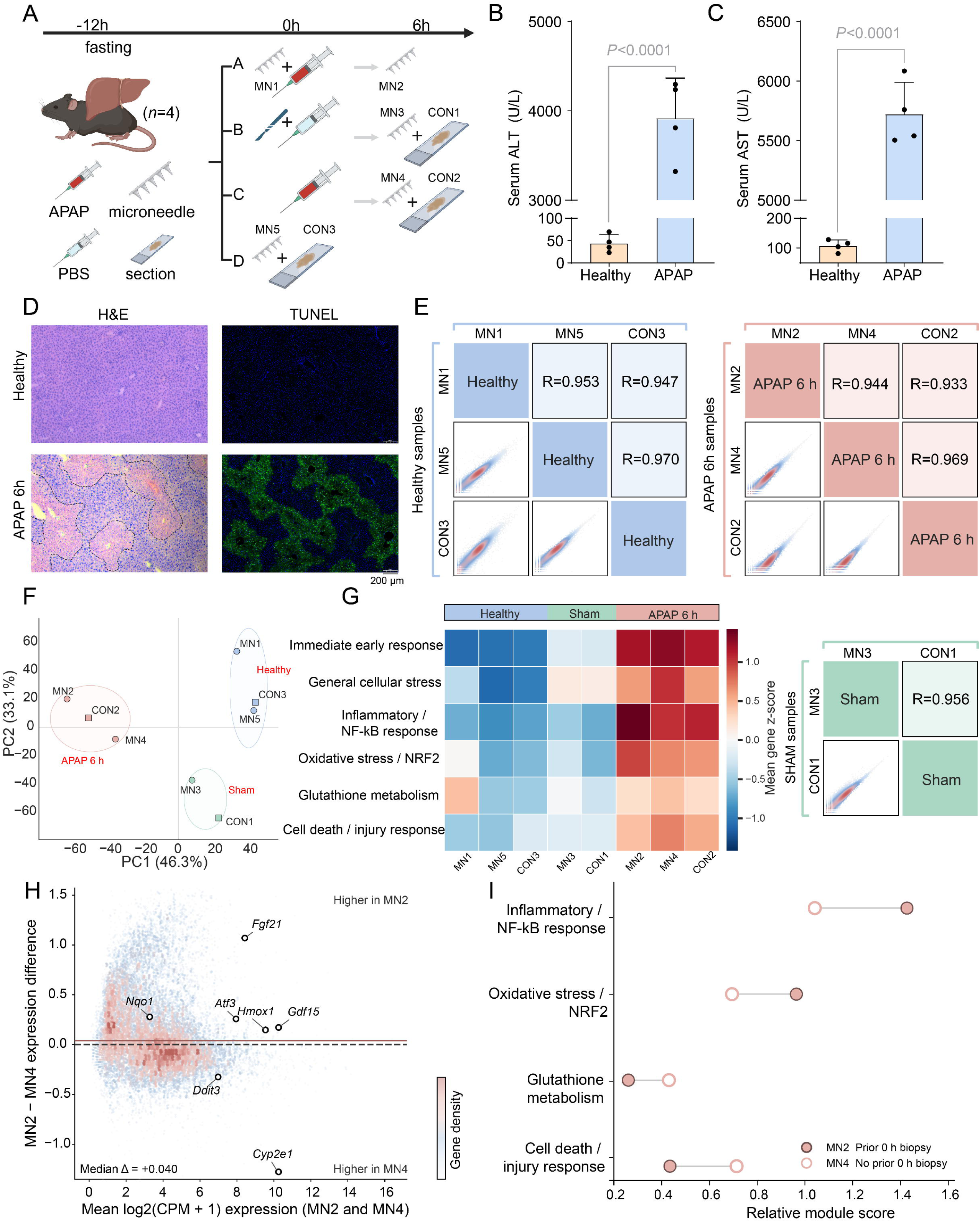
Validation of longitudinal in vivo liver sampling by LANCE-seq. (a) Schematic illustration of the experimental design for evaluating longitudinal in vivo liver sampling. Mouse A underwent baseline microneedle sampling at 0 h (MN1), followed by APAP administration and a second microneedle sampling at 6 h (MN2). Mouse B underwent sham surgery at 0 h without microneedle sampling, received PBS, and was sampled at 6 h to obtain MN3 and CON1. Mouse C received APAP without prior surgery or microneedle sampling and was sampled at 6 h to obtain MN4 and CON2. Mouse D served as an independent healthy control and provided MN5 and CON3. (b, c) Serum alanine aminotransferase (ALT) and aspartate aminotransferase (AST) activities in healthy (*n* = 2) and APAP-treated (*n* = 2) mice, error bar indicates meanß±ßs.d. Statistical significance was assessed using a two-sided unpaired Student’s *t*-test. (d) Representative H&E and TUNEL staining of liver tissues from healthy and APAP 6 h groups, showing APAP-induced hepatic injury and cell death. (e) Pairwise global gene expression correlations among samples within the Healthy, Sham, and APAP 6 h conditions. (f) Principal component analysis (PCA) of the eight transcriptomic samples, showing their global transcriptional relationships across experimental conditions. (g) Comparison of six functional transcriptional modules across Healthy, Sham, and APAP 6 h samples, including immediate early response, general cellular stress, inflammatory/NF-κB signaling, oxidative stress/NRF2 signaling, glutathione metabolism, and cell death/injury. (h) Gene-level comparison of expression differences between MN2, obtained after 0 h surgery and microneedle sampling, and MN4, obtained without a prior 0 h intervention. (i) Comparison of APAP injury-related functional modules between MN2 and MN4.

First, we verified the APAP-induced acute liver injury model through serum biochemistry and histopathology. Compared with the healthy group, serum alanine aminotransferase (ALT) and aspartate aminotransferase (AST) activities were significantly elevated 6 h after APAP treatment (*P* < 0.0001, Fig. 4b, c). Hematoxylin and eosin (H&E) staining revealed obvious hepatocyte damage and necrotic areas in the APAP group, while terminal deoxynucleotidyl transferase dUTP nick-end labeling (TUNEL) staining showed numerous positive signals in the damaged regions, indicating that marked acute liver injury had developed 6 h after APAP treatment (Fig. 4d). In addition, the liver and other tissues after microneedle sampling mainly exhibited histological changes confined to the puncture area, while the surrounding tissue architecture remained largely intact (Supplementary Fig. 3). The distribution of gene and UMI counts detected at each spatial spot across the 8 samples is shown in Supplementary Fig. 4. Further global gene expression correlation analysis revealed that samples within the Healthy, Sham, and APAP 6 h conditions displayed high correlations in their overall expression profiles (Fig. 4e).

On this basis, we compared the overall transcriptional states under three conditions. Principal component analysis (PCA) showed a modest shift of sham-operated samples relative to healthy samples, whereas APAP 6 h samples were more clearly separated from healthy samples (Fig. 4f). Functional module analysis revealed that the Sham condition was primarily characterized by immediate early responses and elevated general cellular stress. In contrast, the APAP 6 h condition exhibited these stress features together with more pronounced transcriptional changes related to inflammation/NF-κB, oxidative stress/NRF2, and cell death/injury, accompanied by alterations in glutathione metabolism (Fig. 4g). These results indicate that surgery and related procedures were associated with a degree of transcriptional stress, whereas APAP treatment was associated with broader transcriptional changes involving oxidative stress, inflammation, and cellular injury.

To further evaluate whether prior 0 h surgery and microneedle sampling systematically affected subsequent APAP responsiveness, we compared MN2, which underwent 0 h surgery and microneedle sampling, with MN4, which received no 0 h intervention. The global gene expression changes in MN2 relative to MN4 did not show a consistent upward or downward shift (Fig. 4h), and the differences in APAP injury-related functional modules between the two also did not exhibit a consistent direction (Fig. 4i). Collectively, under the current experimental conditions, no systematic shift in the overall transcriptional state or major injury programs of the 6 h APAP samples was observed as a result of prior 0 h surgery and microneedle sampling, supporting the feasibility of LANCE-seq for repeated sampling in the same living animal.

### LANCE-seq Reveals Spatiotemporal Transcriptomic Remodeling during APAP-Induced Liver Injury

We further utilized spatial transcriptomic data from the same mouse before and after APAP treatment to analyze liver injury-associated longitudinal changes (Fig. 5a). Differential expression analysis between MN2 and MN1 revealed widespread alterations in the liver transcriptome following APAP treatment, with stress- and injury-related genes such as *Hmox1*, *Ddit3*, *Atf3*, *Fgf21*, and *Gdf15* significantly upregulated, while metabolism-related genes such as Cyp2e1 were downregulated (Fig. 5b). The upregulated genes were primarily enriched in processes related to inflammatory response, neutrophil chemotaxis, cytokine and chemokine signaling, integrated stress response, and endoplasmic reticulum stress (Fig. 5c). Further functional enrichment analysis revealed that upregulated genes were also enriched in signaling pathways such as NF-κB, TNF, IL-17, and MAPK, while downregulated genes were predominantly enriched in pathways related to lipid and cholesterol biosynthesis, mitochondrial function, DNA metabolism and repair, as well as drug metabolism and cytochrome P450-related pathways (Supplementary Fig. 5).

**Fig. 5.**
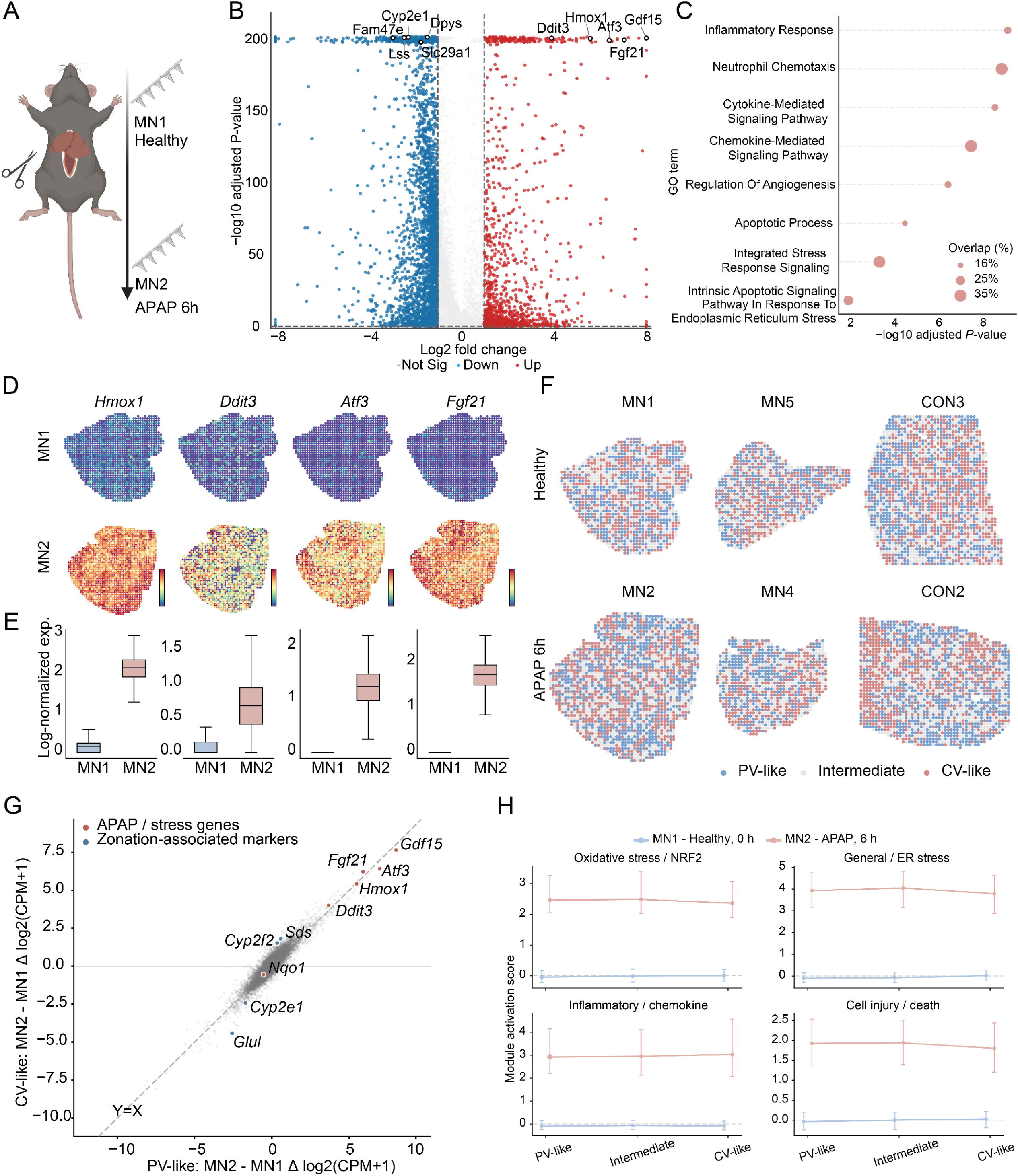
LANCE-seq reveals spatiotemporal transcriptomic remodeling during APAP-induced liver injury. (a) Schematic illustration of longitudinal in vivo sampling from the same mouse before APAP administration (MN1, 0 h) and 6 h after APAP administration (MN2), with the two samplings performed at different liver locations. (b) Volcano plot showing gene expression changes between MN2 and MN1. Representative APAP-responsive genes are labeled. (c) Gene Ontology (GO) enrichment analysis of genes upregulated in MN2 relative to MN1, highlighting inflammatory, stress-response, and cell injury-related biological processes. (d, e) Spatial expression maps (d) and corresponding boxplots of log-normalized expression (e) for representative APAP-responsive genes *Hmox1*, *Ddit3*, *Atf3*, and *Fgf21* in MN1 and MN2. (f) Spatial distribution of relative liver zonation states across Healthy and APAP 6 h samples. Spatial spots were classified as relative portal vein-like (PV-like), intermediate, or relative central vein-like (CV-like) states based on within-sample zonation scores. (g) Comparison of MN2-to-MN1 gene expression changes between relative PV-like and relative CV-like states. The x- and y-axes represent Δlog2(CPM + 1) in PV-like and CV-like states, respectively. Representative APAP-responsive and zonation-associated genes are labeled. (h) Activation of four APAP injury-related transcriptional modules across PV-like, intermediate, and CV-like states in MN1 and MN2, standardized relative to the MN1 baseline. The four modules are oxidative stress/NRF2, general/ER stress, inflammatory/chemokine, and cell injury/death.

We further mapped representative injury-associated genes back to their spatial coordinates. Compared with MN1, *Hmox1*, *Ddit3*, *Atf3*, and *Fgf21* showed higher overall expression in MN2 while retaining pronounced heterogeneity across spatial spots (Fig. 5d, e). Meanwhile, canonical liver zonation-related marker genes also underwent marked changes, with *Glul* and *Cyp2e1* globally decreased in MN2, whereas *Cyp2f2* and *Sds* globally increased, displaying distinct spatial expression patterns (Supplementary Fig. 6). These results indicate that both injury-associated transcriptional responses and liver zonation-related expression features were substantially altered following APAP treatment.

To further analyze the changes in liver zonation-associated transcriptional features after APAP treatment, we referenced previous liver zonation transcriptomic studies and selected genes with relatively stable zonation bias to construct a relative zonation score^29, 30^. Among these, *Rgn*, *Slc1a2*, *Cyp2c29*, and *Gulo* were used as central vein (CV)-associated marker genes, while *Cyp2f2*, *Sds*, *Hal*, and *Pck1* served as portal vein (PV)-associated marker genes. A relative zonation score was constructed within each sample, and spatial spots were classified into relative PV-like, intermediate, and relative CV-like transcriptional states (Fig. 5f). Continuous relative zonation scoring revealed varying degrees of spatial heterogeneity within each sample (Supplementary Fig. 7a). Independent validation was further performed using *Cyp2e1*, *Glul*, *Cyp7a1*, *Ass1*, and *Alb*, which were not used to construct the zonation score. Among these, *Cyp2e1*, *Glul*, and *Cyp7a1* generally showed increased expression from PV-like to CV-like states, whereas *Ass1* and *Alb* exhibited an opposite trend, consistent with the canonical expression direction of liver lobule zonation (Supplementary Fig. 7b).

Subsequently, we established a healthy reference to further evaluate changes in liver zonation-associated transcriptional programs following APAP treatment. Compared with healthy samples MN1 and MN5, the CV-associated transcriptional program scores in APAP-treated MN2 and MN4 were both reduced, whereas the PV-associated transcriptional program did not exhibit a decline of comparable magnitude (Supplementary Fig. 8). Corresponding spatial mapping likewise revealed that the CV-associated program signals in MN2 and MN4 were globally weakened relative to those in healthy samples.

We further compared the distribution of APAP injury-associated transcriptional responses across different relative zonation states. Zonation stratification analysis revealed that transcriptional changes from MN1 to MN2 were largely consistent between the relative PV-like and relative CV-like states, while a subset of genes exhibited zonation-dependent responses of varying magnitude (Fig. 5g). Functional module analysis showed that oxidative stress/NRF2, general and endoplasmic reticulum stress, inflammation/chemokines, and cell injury/death programs were markedly elevated across all three relative zonation states of MN2 compared with the corresponding states of MN1, whereas the differences among the three states within MN2 were relatively small (Fig. 5h). Global spatial autocorrelation analysis further revealed that multiple representative injury-related genes, zonation-associated genes, and functional modules exhibited positive spatial autocorrelation (Supplementary Fig. 9). In addition, supportive analyses performed on other microneedle and planar tissue samples showed that the zone-stratified transcriptional changes from MN1 to MN2 were positively correlated with the corresponding changes from MN5 to MN4 and from CON3 to CON2 (Supplementary Fig. 10). Overall, APAP-related stress and injury transcriptional programs were broadly enhanced across different relative zonation states while retaining a degree of zonation-associated heterogeneity and spatial organization.

## Discussion

In recent years, ST has substantially improved the spatial resolution of gene expression heterogeneity in complex tissues, yet most sequencing-based methods still rely on *ex vivo* tissue processing, making it difficult to directly track longitudinal molecular changes within the same living individual^3, 11–13^. LANCE-seq combines spatial encoding with tissue-penetrating microneedle arrays, demonstrating transcriptome capture and reconstruction of major spatial structures in frozen mouse brain tissue and further enabling repeated spatial transcriptomic sampling at different time points from the same living animal. Existing needle-based transcriptomic methods have separately demonstrated the feasibility of targeted sampling from ex vivo tissues or repeated sampling *in vivo*. LANCE-seq further integrates high-throughput spatial barcoding with minimally invasive in vivo sampling, enhancing the spatial resolution of repeated in vivo sampling and providing a technical foundation for longitudinal spatial transcriptomic analysis of the same animal.

In the APAP-induced acute liver injury model, LANCE-seq captured transcriptional changes consistent with established features of APAP hepatotoxicity. Following APAP overdose, cytochrome P450-mediated metabolic activation generates N-acetyl-p-benzoquinone imine (NAPQI)^31, 32^, which in turn causes glutathione depletion, mitochondrial protein adduct formation, and sustained mitochondrial oxidative stress^33–35^, accompanied by secondary changes including endoplasmic reticulum stress, inflammatory responses, and cellular injury^36–38^. After 6 h of APAP treatment, stress-related genes such as *Hmox1*, *Ddit3*, *Atf3*, and *Fgf21* were markedly elevated, accompanied by the concurrent enhancement of transcriptional programs associated with oxidative stress/NRF2, general and endoplasmic reticulum (ER) stress, inflammation, and cell damage, consistent with the known features of oxidative stress, ER stress, and secondary inflammation and cell injury during APAP-induced hepatotoxicity. Meanwhile, transcriptional changes in the Sham condition were primarily confined to immediate early responses and general cellular stress, whereas the APAP 6 h condition further exhibited more prominent oxidative stress-, inflammation-, and injury-related programs. Further comparison of MN2, which underwent 0 h surgery and microneedle sampling, with MN4, which did not undergo the 0 h procedure, also revealed no consistent-direction global transcriptional shifts or changes in injury programs. Therefore, although the 0 h surgery and microneedle sampling procedures may introduce certain transcriptional perturbations, they did not mask the major injury features observed 6 h after APAP treatment. This provides preliminary support for the use of LANCE-seq in repeated longitudinal sampling from the same living animal.

APAP-induced liver injury also exhibits pronounced zonation dependence within the hepatic lobule^29, 30, 39^. Normal hepatocytes display broad zonated expression patterns along the PV–CV axis, while enzymes involved in APAP metabolic activation, such as CYP2E1, are primarily enriched in the pericentral region surrounding the CV, rendering this area more susceptible to APAP toxicity^30, 36^. Walesky et al. observed pronounced centrilobular necrosis 6 h after APAP treatment, and found that the transcriptional signature of centrilobular hepatocytes was markedly attenuated, with *Cyp2e1*-positive hepatocytes declining from 67% in the healthy state to approximately 5%. During the adaptive and reparative phase following injury, some centrilobular genes were re-expressed outside their original zones, exhibiting pronounced zonated transcriptional reprogramming^40^. In our study, at 6 h after APAP administration, CV-associated marker genes such as *Glul* and *Cyp2e1* were globally downregulated, and the CV-associated transcriptional program was also markedly attenuated. This change is consistent with the attenuation and remodeling of CV-associated transcriptional features observed in previous studies of APAP-induced liver injury, and further demonstrates that LANCE-seq is capable of capturing this spatial injury signature in longitudinal in vivo sampling.

Previous studies have shown that the expression domains of antioxidant stress genes such as *Txnrd1* and *Gclc* can expand across a broader portion of the hepatic lobule following APAP treatment, suggesting that the adaptive stress response after injury has a wider spatial distribution than the necrotic area^40^. Similarly, spatial reconstruction studies based on single-cell transcriptomics have demonstrated that hepatocytes across different hepatic lobular zones can elicit stress responses during the early phase of APAP exposure, yet the extent of these responses and subsequent cellular outcomes vary among zones, with perivenular hepatocytes being more prone to severe injury and cell death^40^. After 6 h of APAP treatment, we similarly observed that oxidative stress, general and endoplasmic reticulum stress, inflammation, and cell damage-related programs were significantly enhanced across different relative zonal states, while the magnitude of changes in some genes still exhibited zonal differences. Therefore, the stress and injury transcriptional programs at 6 h post-APAP treatment displayed a relatively broad spatial distribution while retaining certain zonation-associated differences. The overall attenuation of CV-associated programs further suggested that the basal transcriptional state of this region was more strongly affected.

LANCE-seq currently has several technical limitations that require further improvement. First, compared with two-dimensional planar capture, the reaction environment on the surface of three-dimensional microneedles is more complex, and the proportion of reads with valid spatial barcodes in the MN group is lower than that in the Section group, suggesting that the liquid-phase reaction on the microneedle surface and the subsequent washing process still have room for further optimization. Limited by the microneedle size, the capture resolution of LANCE-seq remains at the level of multicellular populations. Sampling along the tissue depth may also integrate transcripts from different depths, causing some signals to overlap in the final two-dimensional spatial map^12, 21^. The local tissue deformation caused by microneedle puncture, potential molecular diffusion, and the differences between the actual sampling sections of the MN group and the Section group may collectively affect the fidelity of spatial signals as well as the reconstruction accuracy of certain brain region boundaries and two-dimensional spatial profiles. At the current sampling scale, although unsupervised clustering can reflect the transcriptional heterogeneity within samples, it cannot directly resolve the continuous PV–CV structure of the liver lobule through discrete clustering (Supplementary Fig. 11). Furthermore, the current orthogonal spatial encoding relies on manual alignment and sealing between the PDMS microfluidic channels and the microneedle array. To accommodate the existing encoding workflow, the microneedle array is integrated on a standard 75 × 25 mm silicon substrate, and this larger substrate size increases the operational burden during *in vivo* sampling. Although cutting out the central microneedle region after encoding can improve sampling flexibility, the cut microneedle module cannot be readily re-integrated with the existing microfluidic encoding system for repeated functionalization and encoding, substantially increasing the cost per sampling session. Meanwhile, as the needle pitch and channel dimensions are further reduced, the risks of alignment errors and inter-channel fluid leakage will also increase. Therefore, in the future, more compact and reusable spatially barcoded interfaces still need to be developed to further improve the sampling convenience, spatial resolution, and engineering scalability of the platform. Finally, current longitudinal biological validation is primarily based on repeated sampling from the same animal at 0 h and 6 h. Future studies still need to incorporate independent animals, additional sampling time points, and longitudinal replicates to further verify the stability of the observed spatiotemporal transcriptional changes across different individuals and temporal scales. Further optimization of molecular reactions on three-dimensional surfaces, increased microneedle array density, and the development of self-aligned or automated spatial encoding strategies could further improve the spatial resolution and experimental throughput of LANCE-seq.

In summary, this study developed LANCE-seq, a live-tissue spatial transcriptomic sequencing platform based on encoded MNAs. We validated its capability for spatial transcriptome capture and reconstruction of major spatial structures in frozen mouse brain tissues, and achieved repeated sampling at different time points from the same living animal in an APAP-induced acute liver injury model, capturing injury-related transcriptional changes and liver zonation-associated remodeling. LANCE-seq provides a technical strategy for longitudinal spatial transcriptomic studies of drug responses, tissue injury, and disease progression in living tissues.

## Methods

### Animals and Experimental Design

Male C57BL/6J mice (8–10 weeks old) were housed under a standard 12-h light/dark cycle with ad libitum access to food and water. All animal procedures were conducted in accordance with a protocol approved by the Ethics Committee of CUH, Southeast University (Approval No. 20230206002). Mice used for the APAP-induced acute liver injury experiments were fasted for 12 h before treatment. APAP was administered intraperitoneally at 300 mg/kg in prewarmed PBS.

Four mice were used in a longitudinal sampling experiment with complementary controls. Mouse A underwent laparotomy under anesthesia at 0 h, followed by baseline in vivo microneedle sampling (MN1). The abdomen was then closed, APAP was administered, and the animal was allowed to recover. At 6 h after APAP administration, the liver was re-exposed and a second microneedle sample (MN2) was collected from a different site or lobe. Mouse B underwent the same laparotomy and liver exposure at 0 h but without microneedle sampling, followed by abdominal closure and administration of an equivalent volume of PBS. At 6 h, the liver was re-exposed for microneedle sampling (MN3), and liver tissue collected at the terminal endpoint was used to generate the corresponding planar sample CON1. Mouse C received APAP at 0 h without prior laparotomy or microneedle sampling. At 6 h, the liver was exposed for the first time to obtain MN4, followed by terminal tissue collection for CON2. Mouse D served as an independent healthy control and underwent microneedle sampling to obtain MN5, followed by terminal tissue collection for CON3. Mice were euthanized only after completion of their designated terminal sampling procedures.

### Fabrication of Microneedle Arrays

The LANCE-seq platform used silicon microneedle arrays fabricated using standard microfabrication techniques. Briefly, the arrays were fabricated on a standard 75 × 25 mm silicon substrate, with a 50 × 50 microneedle array (2,500 needles in total) positioned in the central region. Each microneedle had a base diameter of 50 μm, a height of 100 μm, and a center-to-center pitch of 200 μm. Following deep reactive ion etching (DRIE) and cleaning procedures, the microneedle arrays were prepared for subsequent surface functionalization.

### Microfluidic Device Design and Fabrication

A customized PDMS flow cell was designed for the orthogonal spatial encoding of the microneedle arrays^26, 27^. The microfluidic device consisted of parallel microchannels with a channel width and inter-channel spacing of 100 μm, yielding a 200 μm periodicity that matched the center-to-center pitch of the microneedle array. The PDMS slabs were fabricated using conventional soft lithography. A PDMS prepolymer mixture with a base-to-curing-agent ratio of 10:1 was poured onto an SU-8 master mold and cured at 80 °C for 2 h.

### Preparation of 3D Dendrimeric Arrays and 2D Functionalized Slides

The silicon microneedle arrays were functionalized with a three-dimensional PAMAM dendrimer layer using the surface-chemistry strategy previously established for Decoder-seq^41^ to increase the density of reactive sites. Briefly, the hydroxylated silicon surface was functionalized with γ-Glycidoxypropyltrimethoxysilane (GOPTS) to introduce epoxy groups, followed by coupling of generation-4 PAMAM dendrimers through a ring-opening reaction. The resulting amine-rich dendrimeric surface was subsequently used for disuccinimidyl suberate (DSS)-mediated immobilization of amino-modified Read 1 probes. For preparation of the 2D planar control slides, glass slides were cleaned and activated with piranha solution for 2 h, followed by treatment with 5% 3-aminopropyltriethoxysilane in anhydrous ethanol for an additional 2 h. After being thoroughly washed and dried, the 2D slides were baked at 110 °C for 1 h.

### Generation and Orthogonal Spatial Encoding of LANCE-seq Chips

The spatial barcoding procedure was identical for the microneedle arrays and planar glass slides. To initiate spatial barcoding, 40 μM amino-modified Read 1 probes were covalently conjugated to the activated surfaces via DSS-mediated crosslinking. After incubation, the substrates were blocked, sequentially washed, and dried. The oligonucleotide sequences used for surface functionalization, spatial barcoding, reverse transcription, amplification, indexing, and fluorescence validation are provided in Supplementary Table 1.

To generate surfaces with spatially defined DNA barcodes, we employed two different customized PDMS microfluidic devices. First, Chip A was precisely aligned with the rows of the array under a bright-field microscope and held firmly with customized acrylic clamps to prevent liquid leakage. A ligation mixture containing 10 μM of one of the 50 distinct Barcode A probes pre-annealed to its complementary ligation linker and 40 U/μL T4 DNA ligase was prepared. A unique Barcode A mixture was introduced into each parallel inlet of Chip A. After incubation at room temperature for 2 h in an enclosed humid chamber, the barcode solutions were removed, and the channels were continuously rinsed with wash buffer. The acrylic clamps were then removed, and Chip A was detached from the array. The array was sequentially washed with 2× saline–sodium citrate (SSC) containing 0.1% sodium dodecyl sulfate (SDS) and with 0.2× SSC, followed by a rinse with Milli-Q water.

After the array was dried under a stream of clean compressed air, Chip B, whose microchannels were oriented perpendicular to those of Chip A, was aligned with and attached to the dried array. Similarly, a ligation mixture containing 10 μM of one of the 50 distinct Barcode B probes, the corresponding linker, and 40 U/μL T4 DNA ligase was introduced into each inlet, with a unique Barcode B assigned to each channel. The ligation reaction was performed under the same conditions for 2 h, after which the reaction mixture was aspirated and the channels were washed. Following the detachment of Chip B, the spatially barcoded region was incubated in 80 mM KOH for 30 s to remove the complementary linkers. Finally, the completed chips were rinsed with water, dried under a stream of clean air, and stored under inert, light-protected conditions at room temperature until use. To verify spatial encoding and probe uniformity, the chips were hybridized with Cy3-labeled poly(A) probes and imaged by fluorescence microscopy.

### Tissue Collection and *In Situ* Sampling

For *ex vivo* brain tissue analysis, fresh mouse brains were rapidly harvested and snap-frozen in liquid nitrogen. Frozen brains were mounted on a specimen chuck, and LANCE-seq microneedle arrays were pressed onto the coronal tissue surface to achieve tissue penetration and in situ molecular capture.

For *in vivo* liver sampling, mice were anesthetized and the liver was surgically exposed according to the experimental design described above. LANCE-seq microneedle arrays were directly applied to the exposed liver surface for in situ transcriptomic sampling. For animals undergoing longitudinal sampling, the abdomen was closed after the 0 h procedure and the animals were allowed to recover until the scheduled 6 h sampling. Mice were euthanized only after completion of the designated terminal sampling procedure. Liver tissues were then collected, embedded in optimal cutting temperature (OCT) compound, and cryosectioned at a thickness of 10 μm for planar spatial transcriptomic analysis, histological evaluation, and other downstream assays.

### Serum Biochemical Analysis

Whole blood was collected from mice at the designated experimental endpoints, and serum was isolated for biochemical analysis. Serum ALT and AST activities were measured using commercial ALT/GPT (C009-3-2) and AST/GOT (C010-3-2) assay kits from Nanjing Jiancheng Bioengineering Institute (Nanjing, China) according to the manufacturer’s instructions. Enzyme activities were determined based on changes in absorbance at 340 nm.

### Histological Staining and Imaging

OCT-embedded liver and other tissue samples were cryosectioned for histological evaluation. For H&E staining, tissue sections were fixed and stained with hematoxylin for 2 min, followed by bluing for 20 s and counterstaining with eosin for 1 min. The stained sections were subsequently washed, mounted, and imaged by bright-field microscopy. For evaluation of APAP-induced liver injury, OCT-embedded frozen liver tissues were submitted to Wuhan Servicebio Technology Co., Ltd. for TUNEL staining. DNA fragmentation was detected using a commercial TUNEL assay kit, nuclei were counterstained with 4′,6-diamidino-2-phenylindole (DAPI), and the sections were imaged by fluorescence microscopy.

### Tissue Permeabilization and Reverse Transcription

Post-sampling microneedle arrays and planar slides bearing tissue sections were incubated on a heating block at 37 °C for 1 min. The tissues were then fixed in pre-chilled methanol (−20 °C) for 30 min and dried with isopropanol at room temperature for 1 min. To release intracellular mRNA, 100 μL of 0.1% (w/v) pepsin in 0.1 M HCl was applied to the reaction area and incubated at 37 °C for 15 min (13 min for liver tissue). After permeabilization, the microneedle arrays and planar slides were washed three times with 0.1× SSC.

For in situ reverse transcription, a 75 μL reverse transcription (RT) mixture was prepared containing 5% PEG 8000, 1× RT buffer, 10 U/μL Maxima H Minus Reverse Transcriptase, 2 U/μL RNase inhibitor, 0.5 mM dNTP mix, 2.5 μM template-switching oligo (TSO), 1 mM GTP, and RNase-free water. The RT mixture was added to the reaction region and incubated at 42 °C for 2 h to synthesize the first-strand cDNA. After the reaction, the chips were washed three times with 0.1× SSC buffer. To degrade the hybridized mRNA template, 100 μL of 80 mM KOH was added and incubated at room temperature for 5 min, followed by three additional washes with 0.1× SSC.

### Second-Strand Synthesis, Library Construction, and Sequencing

Second-strand cDNA synthesis was performed directly on the chip using a 37.5 μL reaction mixture containing 4 mM MgSO4, 1× isothermal amplification buffer, 0.32 U/μL Bst 2.0 DNA polymerase (NEB, M0537S), 1 mM dNTP mix, 2.5 μM second-strand primer, and RNase-free water. The reaction was incubated at 65 °C for 15 min. Subsequently, the chips were washed once with 0.1× SSC and twice with Elution Buffer (EB). To cleave and elute the spatially barcoded double-stranded cDNA constructs from the array surface, 40 μL of 80 mM KOH was added and incubated at room temperature for 10 min. The eluate was then transferred to a PCR tube and neutralized with 6 μL of 1 M Tris buffer (pH 7.0).

For cDNA amplification, 50 μL of KAPA Hotstart ReadyMix and 2 μL each of 20 μM amplification primers (Amp-F and Amp-R) were added to the neutralized cDNA solution. PCR thermal cycling was performed as follows: initial denaturation at 98 °C for 3 min; 14 cycles of 98 °C for 20 s, 65 °C for 30 s, and 72 °C for 2 min; followed by a final extension at 72 °C for 5 min. The amplified cDNA was purified using magnetic beads at a 0.7× bead-to-sample ratio. cDNA concentration was measured using a Qubit 4.0 Fluorometer, and fragment-size analysis was performed at the Nanjing Jiangbei New Area Biomedical Public Service Platform. Finally, 5 ng of purified cDNA was subjected to fragmentation, adapter ligation, and sequencing-library construction using the TruePrep DNA Library Prep Kit V2 (Vazyme, TD512) according to the manufacturer’s protocol. The resulting libraries were purified, quantified, and sequenced on an Illumina NovaSeq 6000 platform with 150-bp paired-end reads (PE150).

### Sequencing Data Processing

Raw paired-end sequencing reads were first preprocessed using custom Python scripts. According to the LANCE-seq capture-probe architecture, two 8-bp spatial barcode sequences and a 12-bp unique molecular identifier (UMI) were extracted from Read 1 and reconstructed into a 16-bp spatial barcode followed by the 12-bp UMI. The first 30 bp of Read 2, containing adapter- and template-switching-related sequences, were trimmed before transcript alignment. Processed reads were aligned to the mouse GRCm39 reference genome using STARsolo (STAR v2.7.11)^42^. A predefined whitelist corresponding to the 50 × 50 combinatorial barcode array was used for spatial barcode assignment, UMI deduplication, and gene-level quantification. Valid Barcode A–Barcode B combinations were mapped to their corresponding two-dimensional positions on the microneedle or planar array, generating a raw gene-by-spatial-spot count matrix for the 2,500 predefined spatial positions.

For planar tissue samples (CON), the spatial barcode array was registered to the corresponding H&E image, and tissue-covered spots were distinguished from background spots based on the tissue boundary. Spots outside the target tissue region were removed before downstream analysis. For microneedle samples (MN), no complete planar tissue section directly corresponded to the sampled array. Valid sampling regions were therefore identified from the raw spatial expression matrix together with spot-level gene and UMI detection metrics, and low-signal background spots were removed. The resulting spatial count matrices were used for downstream quality control, normalization, and transcriptomic analysis.

### Data Preprocessing and Clustering

Downstream analysis of spatial transcriptomic data was primarily performed using Scanpy^43^. Spots with fewer than 50 observed genes (UMI > 2) were filtered out. Genes detected in fewer than 10 spatial spots were discarded. Additional low-quality spots were excluded using sample-specific lower and upper total UMI-count thresholds and a mitochondrial transcript fraction cutoff of 20%. The filtered expression matrices were normalized to 10,000 total counts per spatial spot and log-transformed using log1p for subsequent dimensionality reduction, clustering and gene-expression analyses.

For joint analysis of multiple samples, PCA was performed on highly variable genes, followed by Harmony v0.0.9 integration to reduce technical variation among samples^44^. A transcriptome-based neighborhood graph was subsequently constructed, followed by uniform manifold approximation and projection (UMAP) visualization and unsupervised clustering using the Leiden algorithm^45^. Representative marker genes for each cluster were identified using the Wilcoxon rank-sum test. Cluster assignments and gene-expression patterns were mapped back to the original spatial coordinates for spatial visualization. For independent within-sample clustering of MN1 and MN2, each sample was processed and clustered separately without cross-sample batch correction.

### Brain Region Annotation and Cell-type Deconvolution

Mouse brain transcriptional clusters were annotated according to their spatial distributions and representative region-associated marker genes, with spatial expression patterns of selected markers further validated using in situ hybridization data from the ABA^28^. Cell-type deconvolution was performed using cell2location (v0.1.3)^46^, with the publicly available mouse whole-brain single-nucleus RNA-sequencing dataset E-MTAB-11115 as the reference. Cell-type-specific expression signatures derived from the reference dataset were matched to the LANCE-seq mouse brain spatial expression matrix to estimate the relative abundance of different cell types at each spatial spot. The inferred abundances were mapped back to the original spatial coordinates, and 24 representative neuronal and glial subpopulations were selected for visualization in Fig. 3.

### Transcriptomic and Functional Analysis

For global transcriptomic comparisons, raw counts from all spatial spots within each sample were aggregated by gene, converted to counts per million (CPM), and transformed as log2(CPM + 1). Overall transcriptional similarity between samples was evaluated using Pearson correlation analysis, and principal component analysis (PCA) was used to assess global transcriptional relationships among experimental conditions. Functional transcriptional programs in Fig. 4 were evaluated using predefined gene sets representing immediate early response, general cellular stress, inflammatory/NF-κB signaling, oxidative stress/NRF2 signaling, glutathione metabolism, and cell death/injury, as listed in Supplementary Table 2. For each functional module, individual genes were z-score standardized across the eight samples, and the mean standardized expression of the available genes within each module was calculated as the module score. Gene-level differences between MN2 and MN4 were calculated from their corresponding sample-level log2(CPM + 1) expression profiles and used for descriptive comparison of the two APAP 6 h samples.

Differential-expression analysis between MN1 and MN2 was performed as a within-animal, spot-level comparison using the normalized expression matrix. Differentially expressed genes were identified using the Wilcoxon rank-sum test with thresholds of adjusted *P* < 0.05 and |log2(fold change)| > 1. Upregulated and downregulated genes were subjected separately to Gene Ontology Biological Process (GO-BP) and Kyoto Encyclopedia of Genes and Genomes (KEGG) pathway enrichment analyses using GSEApy^47^. Enrichment *P* values were adjusted for multiple testing using the Benjamini–Hochberg false-discovery-rate procedure.

### Liver Zonation and Spatial Analysis

Liver zonation analysis was performed with reference to previous studies^29, 30^. *Rgn*, *Slc1a2*, *Cyp2c29*, and *Gulo* were used as the CV-associated core gene set, whereas *Cyp2f2*, *Sds*, *Hal*, and *Pck1* were used as the PV-associated core gene set. Log-normalized expression of each core gene was z-score standardized within each sample, and the relative zonation score was calculated as the mean standardized expression of the CV-associated genes minus that of the PV-associated genes. Spatial spots were ranked by zonation score and divided into three equally sized groups representing relative PV-like, intermediate, and relative CV-like states. *Cyp2e1*, *Glul*, *Cyp7a1*, *Ass1*, and *Alb*, which were not included in score construction, were used to validate the relative zonation ordering.

To assess changes in zonation-associated transcriptional programs following APAP treatment, healthy microneedle samples MN1 and MN5 were equally weighted to define gene-specific healthy-reference means and variances, with the variance accounting for both within-sample and between-sample variation. Expression was standardized using these fixed reference parameters, and CV- and PV-associated program scores were calculated as the mean standardized expression of the corresponding core genes.

For zone-stratified analyses, gene expression was aggregated separately within the PV-like, intermediate, and CV-like states of MN1 and MN2. Gene-expression changes were expressed as Δlog2(CPM + 1), calculated as MN2 minus MN1. For the APAP-related module analysis, the mean and standard deviation of each module gene were calculated across MN1 spatial spots and used to standardize the corresponding expression values in both MN1 and MN2. The mean standardized expression of genes within each module was then calculated for each spatial spot as the module activation score. The four functional programs analyzed were oxidative stress/NRF2, general/ER stress, inflammatory/chemokine, and cell injury/death, as listed in Supplementary Table 2. Zone-stratified transcriptional changes in additional independent samples were compared with the MN1-to-MN2 response using Spearman correlation. Global spatial autocorrelation of representative genes, the zonation score, and functional modules was assessed using Global Moran’s I^48^ based on symmetric binary six-nearest-neighbor spatial weights. Significance was evaluated using 999 permutations, with multiple-testing correction performed using the Benjamini–Hochberg method.

### Statistics and Reproducibility

Unless otherwise stated, P < 0.05 was considered statistically significant, and multiple testing was corrected using the Benjamini–Hochberg method. Pearson correlation coefficients were used to evaluate overall similarity between sample-level gene-expression profiles, whereas Spearman correlation coefficients were used for zonation-score validation and concordance analyses of zone-stratified transcriptional changes. Differential-expression analyses were performed using the Wilcoxon rank-sum test. Serum ALT and AST comparisons between healthy and APAP-treated mice were performed using a two-sided unpaired Student’s t-test.

For boxplots, boxes represent the first and third quartiles, center lines indicate the median, and whiskers extend to the largest and smallest values within 1.5 times the interquartile range. Fluorescence intensity measurements were quantified using ImageJ. Statistical analyses were performed in Python within Jupyter Notebook and in GraphPad Prism. Data processing and visualization used NumPy, pandas, Matplotlib, seaborn, and related Python packages. All code used for the analyses is available as described in the Code Availability section.

## Author contributions

W.W. and H.W. designed the experiments, performed the experimental work and developed the analysis code. W.W. analyzed the data and wrote the paper. L.H. prepared the figures. K.D. contributed to the experimental design. L.C. and C.G. contributed to technical discussions and provided advice. K.Y. provided experimental guidance and coordinated the study. X.Z. acquired funding and supported the project. All authors reviewed and approved the paper.

## Data availability

The raw sequencing data generated in this study have been deposited in the Genome Sequence Archive (GSA) in the National Genomics Data Center under accession code CRA079844. Processed spatial transcriptomic data, including expression matrices, spot metadata and spatial coordinates in AnnData format, are available at Zenodo under DOI https://doi.org/10.5281/zenodo.22121908. The mouse whole-brain snRNA-seq dataset used for cell-type deconvolution was obtained from ArrayExpress under accession E-MTAB-11115. Allen Brain Atlas *in situ* hybridization data were obtained from the Allen Brain Atlas.

## Code availability

Custom FASTQ preprocessing scripts and key analysis code generated in this study are available via GitHub at https://github.com/Wong-WJ/LANCE-seq.

