## Supplementary information for "LANCE-seq: Live-tissue Analysis via Needle-encoded Capture Extraction for Spatial Transcriptomics"

**Supplementary Materials**

**Wenjia Wang^1#^, Handong Wang^1#^, Liyong He^2^,** **Kaitong Dang^3^, Liuxi Chu^4^, Chen Gao^5^, Kaiqiang Ye^1,^*, Xiangwei Zhao^1,^***

*^1^State Key Laboratory of Digital Medical Engineering, School of Biological Science & Medical Engineering, Southeast University, Nanjing, Jiangsu 210096, China*

*^2^Medical Research Center, Jiangsu Provincial Medical Key Discipline (Laboratory) Cultivation Unit of Immunology, Nantong First People’s Hospital, Southeast University, Nantong,* *Jiangsu 226006, China*

*^3^Frontier Institute of Science and Technology, Xi' an Jiaotong University, Xi' an, Shaanxi 710049, China*

*^4^The First Affiliated Hospital of Wenzhou Medical University, Wenzhou, Zhejiang 325000, China*

*^5^Department of Modern Mechanics, University of Science and Technology of China, Hefei, Anhui 230027, China.*

**^#^**These authors contributed equally

^*^Corresponding author:

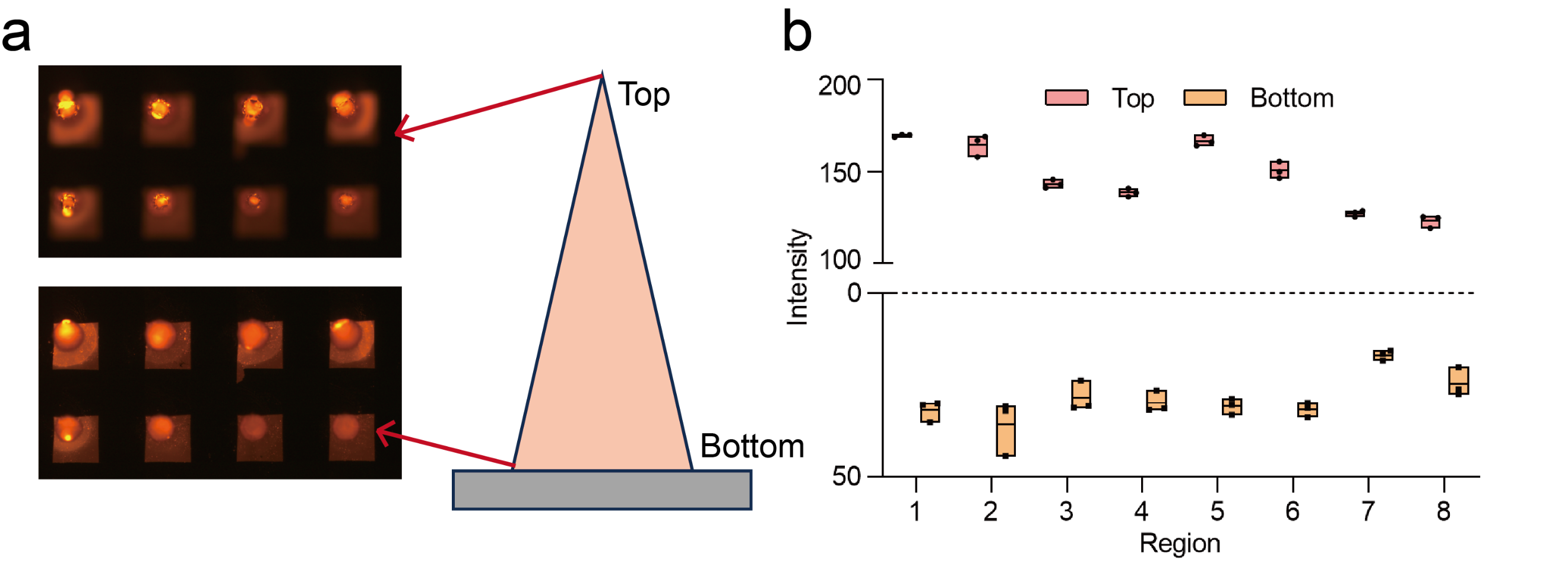

**Supplementary Fig. 1 Uniformity of spatially encoded probes on 3D microneedles.** (a) Fluorescence imaging of the top (tip) and bottom (base) focal planes of the microneedle array. (b) Quantification of fluorescence intensity across different positions of the array. The microneedle tip plane showed higher fluorescence intensity than the bottom plane.

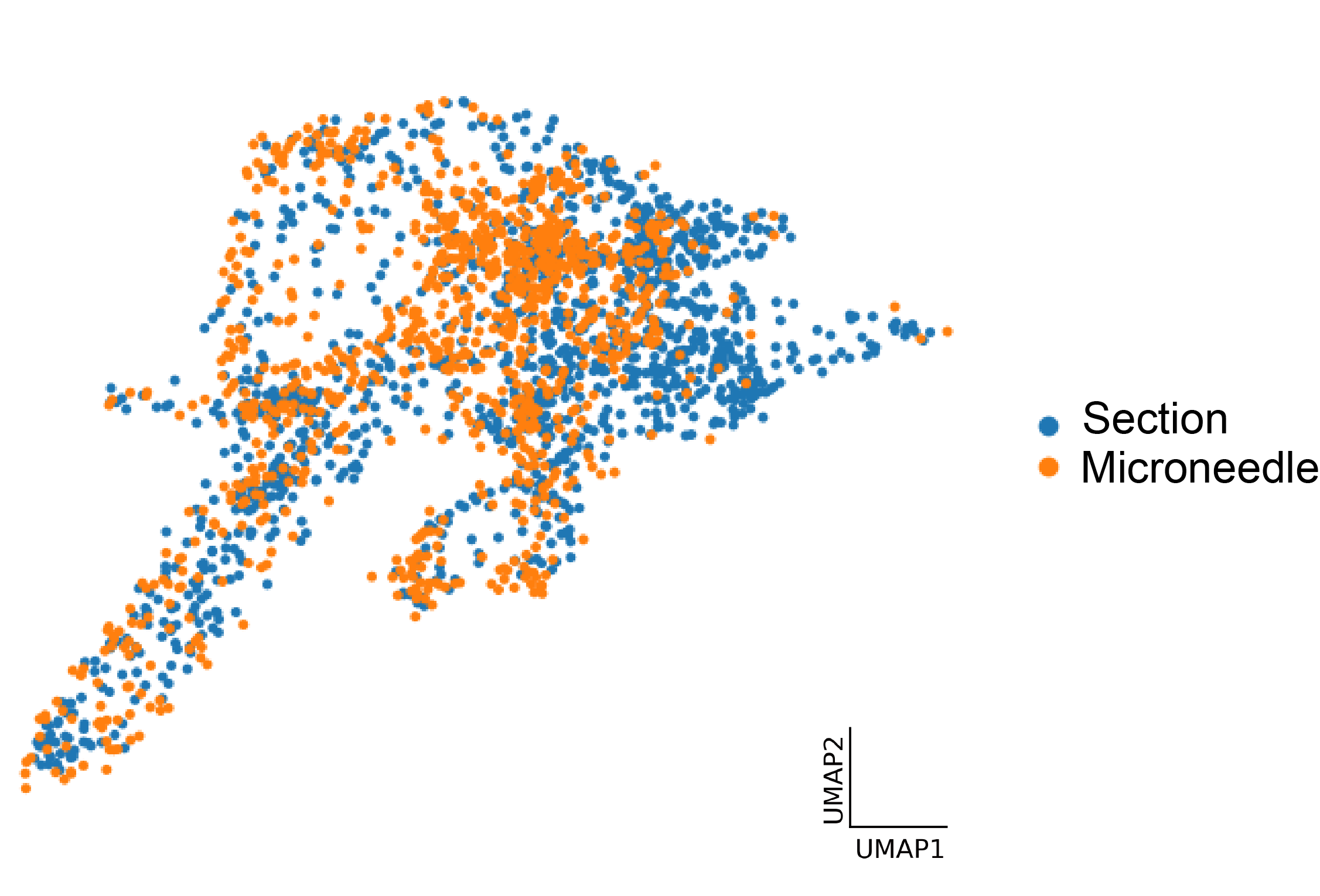

**Supplementary Fig. 2 UMAP visualization of frozen mouse brain spatial transcriptomic data by capture method.** Spatial spots from the planar Section and Microneedle datasets are projected onto the shared UMAP space and colored according to the corresponding capture method.

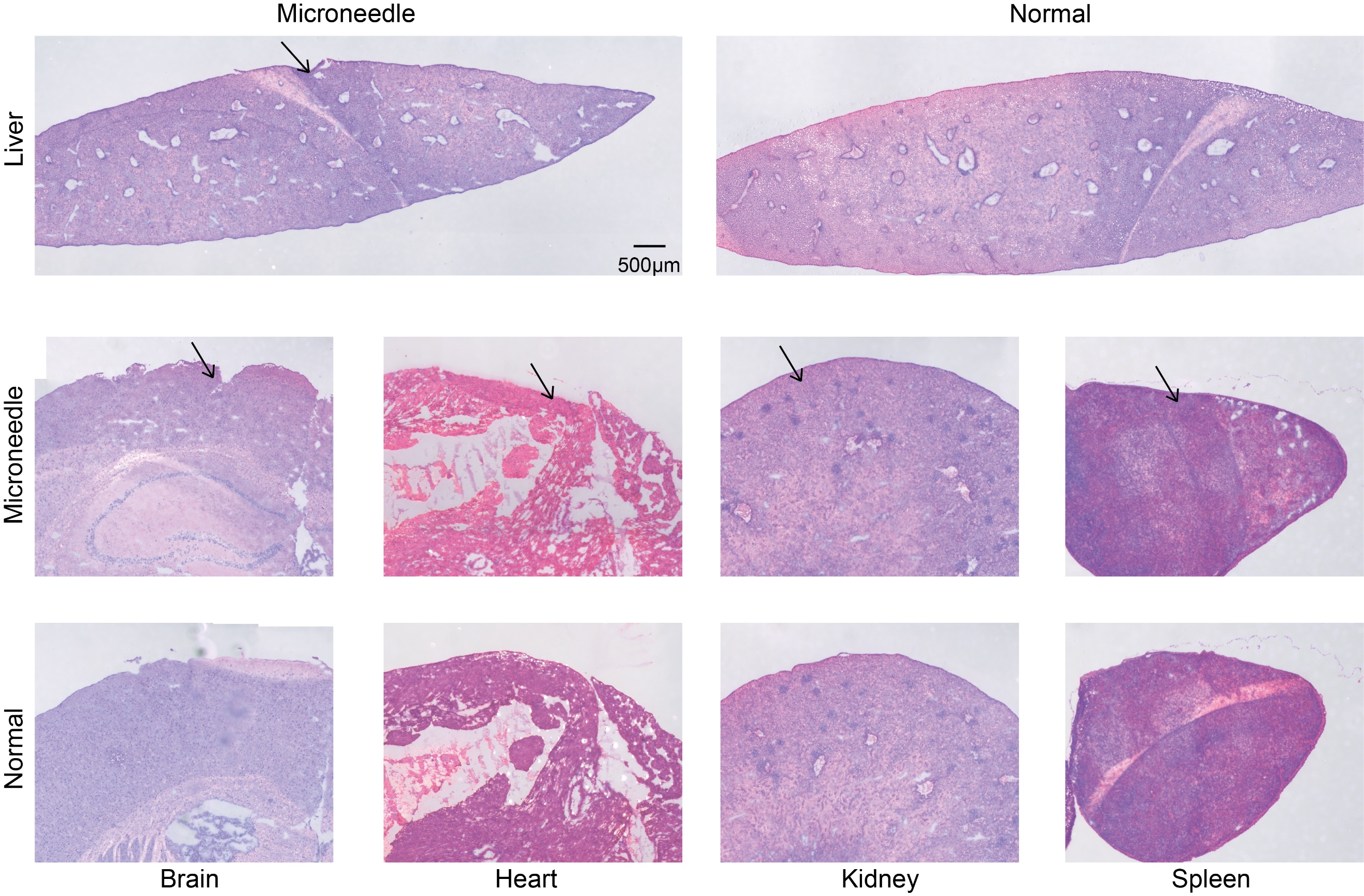

**Supplementary Fig. 3 Histological evaluation of tissue morphology following microneedle sampling.** Representative H&E-stained images of microneedle-sampled tissues and corresponding unsampled controls are shown. The upper panels show liver tissues following microneedle sampling and unsampled control liver tissues, with arrows indicating visible local puncture sites. The lower panels show representative brain, heart, kidney, and spleen tissues following microneedle sampling together with the corresponding unsampled controls. Arrows indicate identifiable local puncture tracks. The overall tissue architecture surrounding the sampled regions remained largely preserved. Scale bar, 500 μm for the liver images.

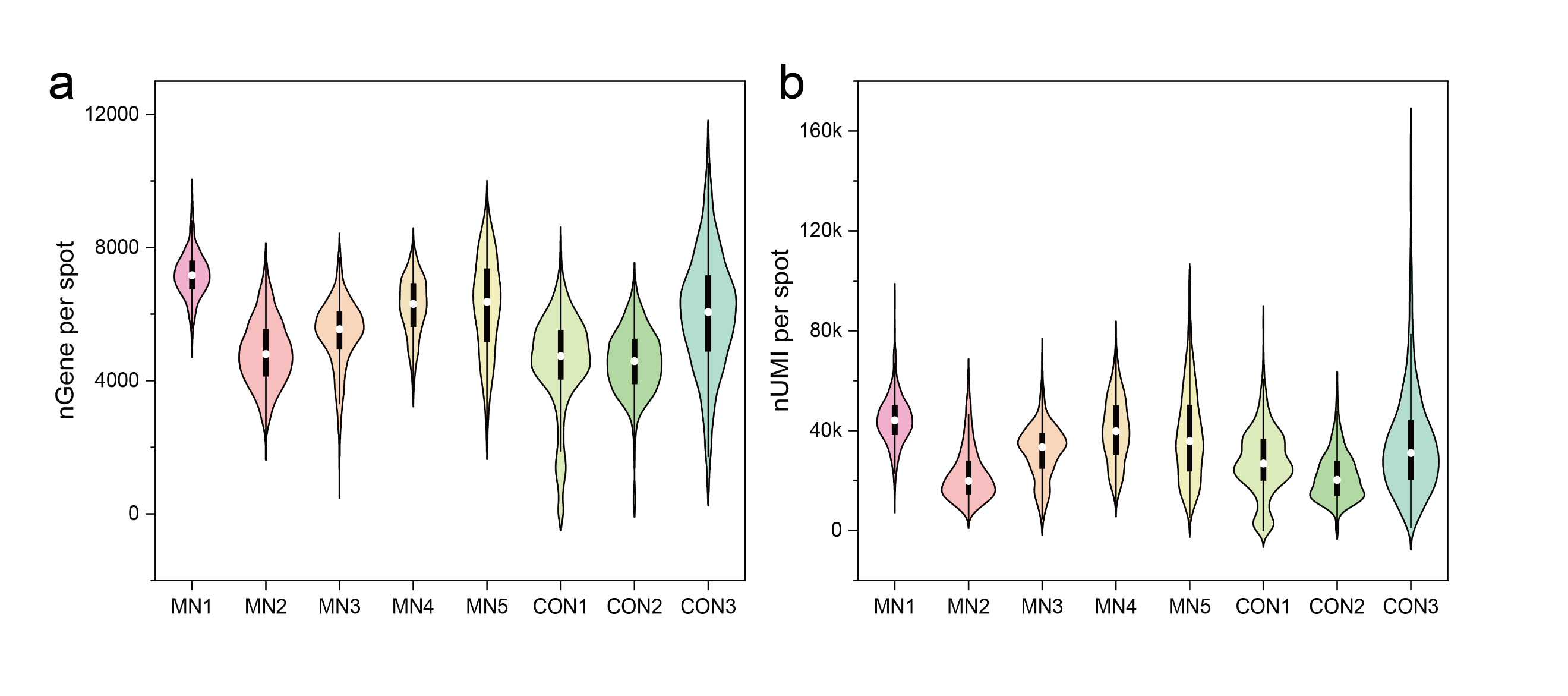

**Supplementary Fig. 4 Distributions of detected genes and UMIs across liver spatial transcriptomic samples.** (a) Distribution of the number of detected genes per spatial spot in MN1–MN5 and CON1–CON3. (b) Distribution of UMI counts per spatial spot in the corresponding samples.

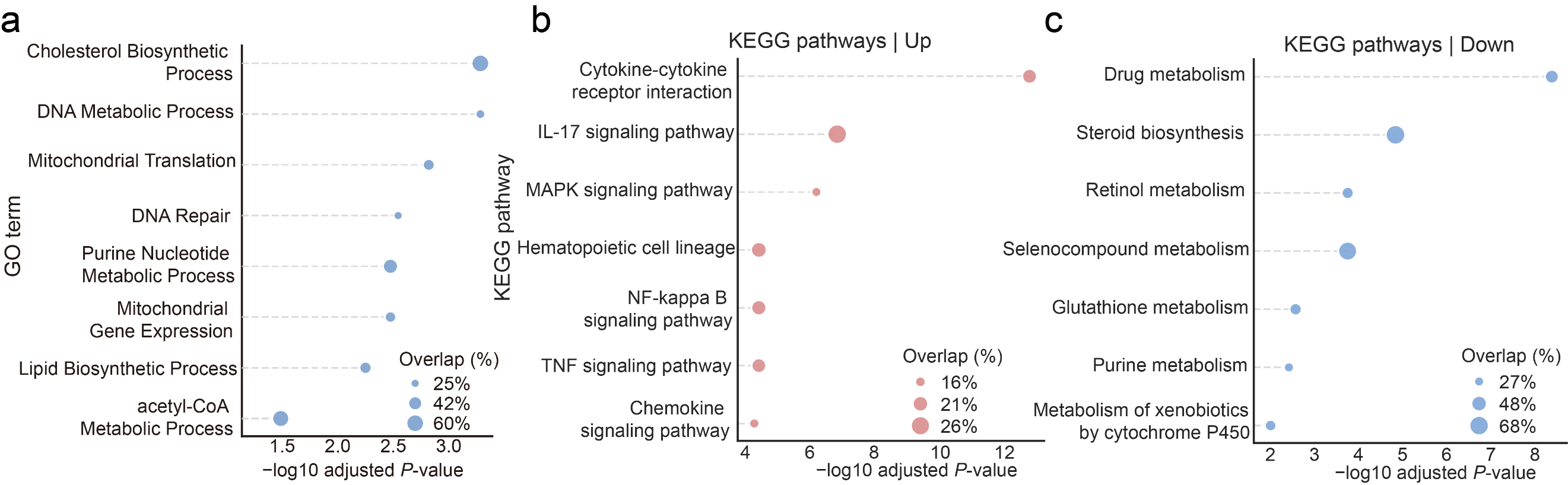

**Supplementary Fig. 5 Functional enrichment analysis of differentially expressed genes between MN2 and MN1.** (a) Gene Ontology Biological Process (GO-BP) enrichment analysis of downregulated genes. (b, c) KEGG pathway enrichment analysis of upregulated (b) and downregulated (c) genes. Bubble size represents the percentage of differentially expressed genes overlapping with each term or pathway.

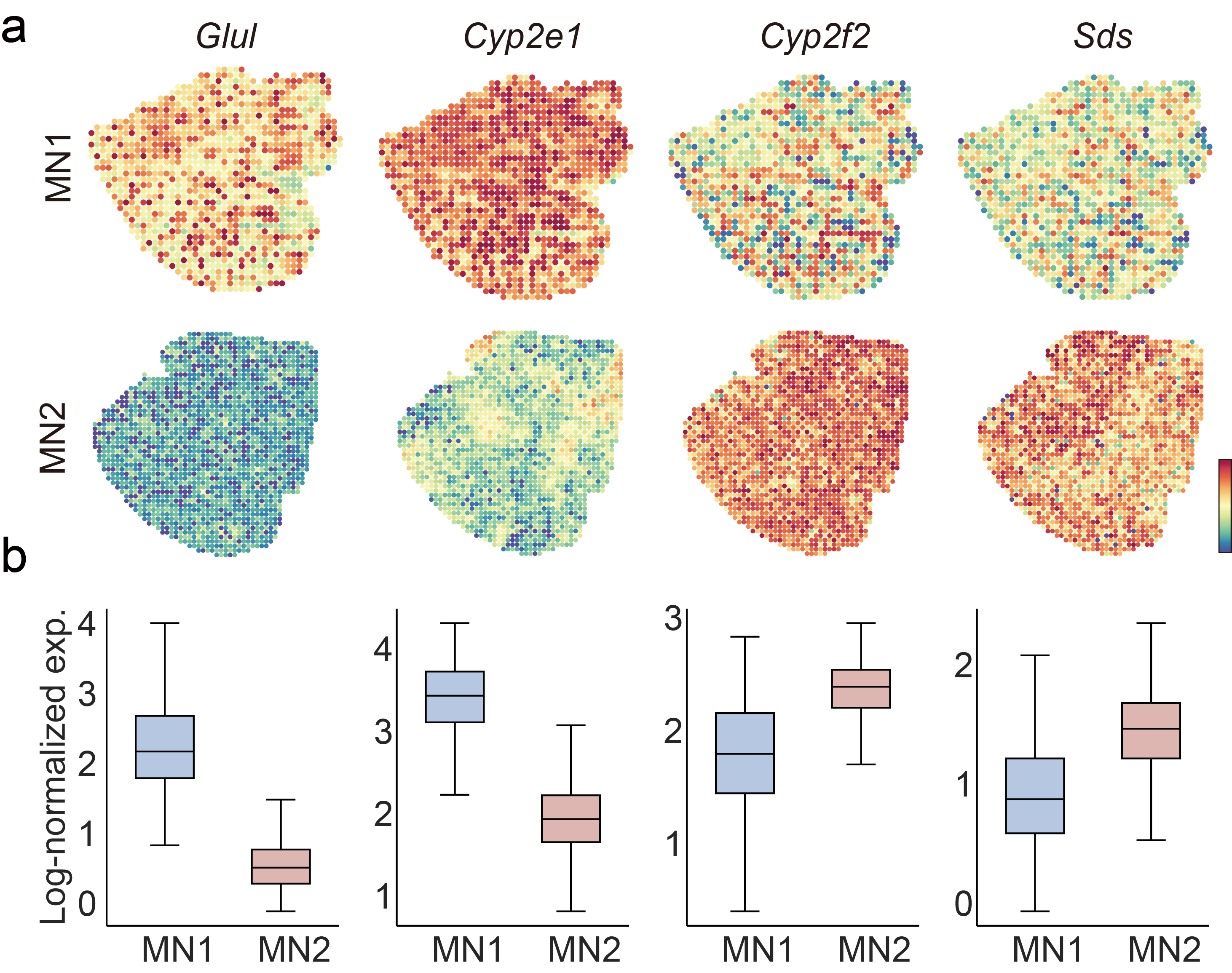

**Supplementary Fig. 6 Spatial expression patterns of representative liver zonation-associated genes in MN1 and MN2.** (a) Spatial expression maps of the central vein (CV)-associated genes *Glul* and *Cyp2e1* and the portal vein (PV)-associated genes *Cyp2f2* and *Sds* in MN1 and MN2. (b) Corresponding boxplots showing the log-normalized expression of *Glul*, *Cyp2e1*, *Cyp2f2*, and *Sds* across spatial spots in MN1 and MN2.

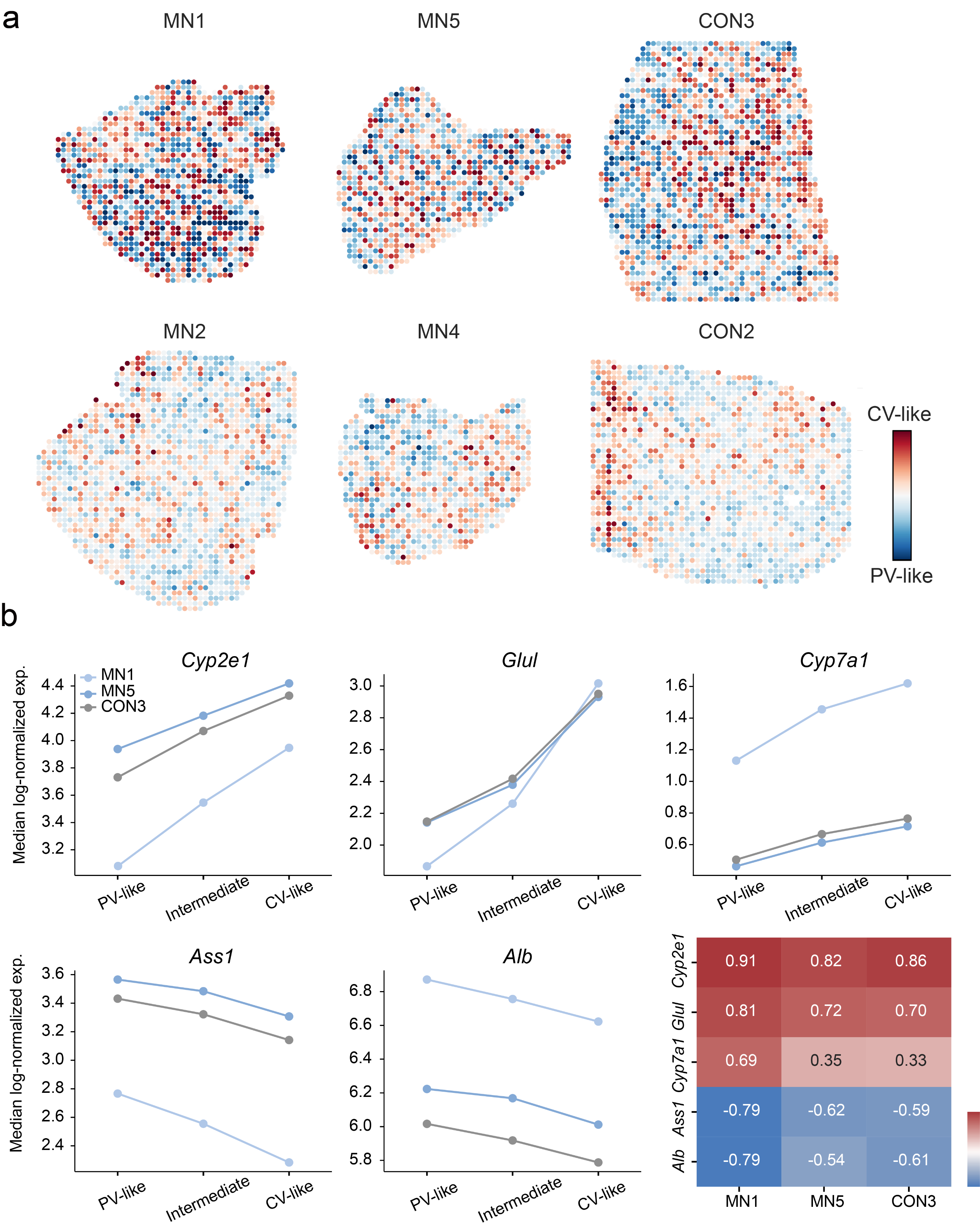

**Supplementary Fig. 7 Continuous relative zonation maps and independent validation of within-sample zonation states.** (a) Spatial distribution of continuous relative zonation scores across Healthy (MN1, MN5, and CON3) and APAP 6 h (MN2, MN4, and CON2) samples. Higher scores indicate more CV-like transcriptional states, whereas lower scores indicate more PV-like states. (b) Independent validation of the relative zonation states in healthy samples using genes not included in the construction of the zonation score. Median log-normalized expression of *Cyp2e1*, *Glul*, *Cyp7a1*, *Ass1*, and *Alb* is shown across PV-like, intermediate, and CV-like states in MN1, MN5, and CON3. The heatmap summarizes the correlation between each held-out marker and the continuous zonation score across the three healthy samples.

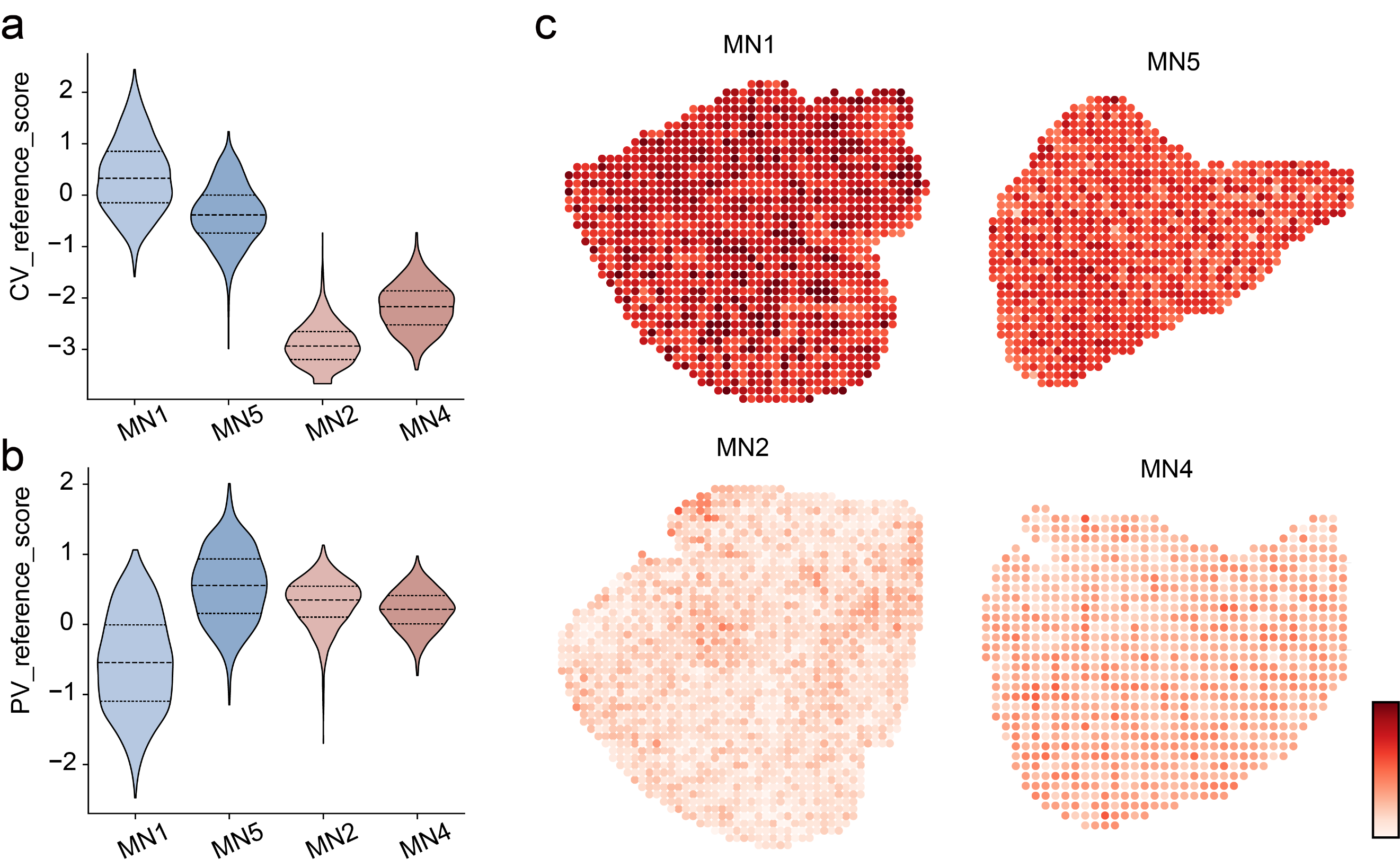

**Supplementary Fig. 8 Healthy-reference analysis of liver zonation program remodeling after APAP treatment.** (a, b) Distributions of healthy-reference CV-associated (a) and PV-associated (b) program scores in MN1, MN5, MN2, and MN4. (c) Spatial maps of the healthy-reference CV-associated transcriptional program score in the same samples.

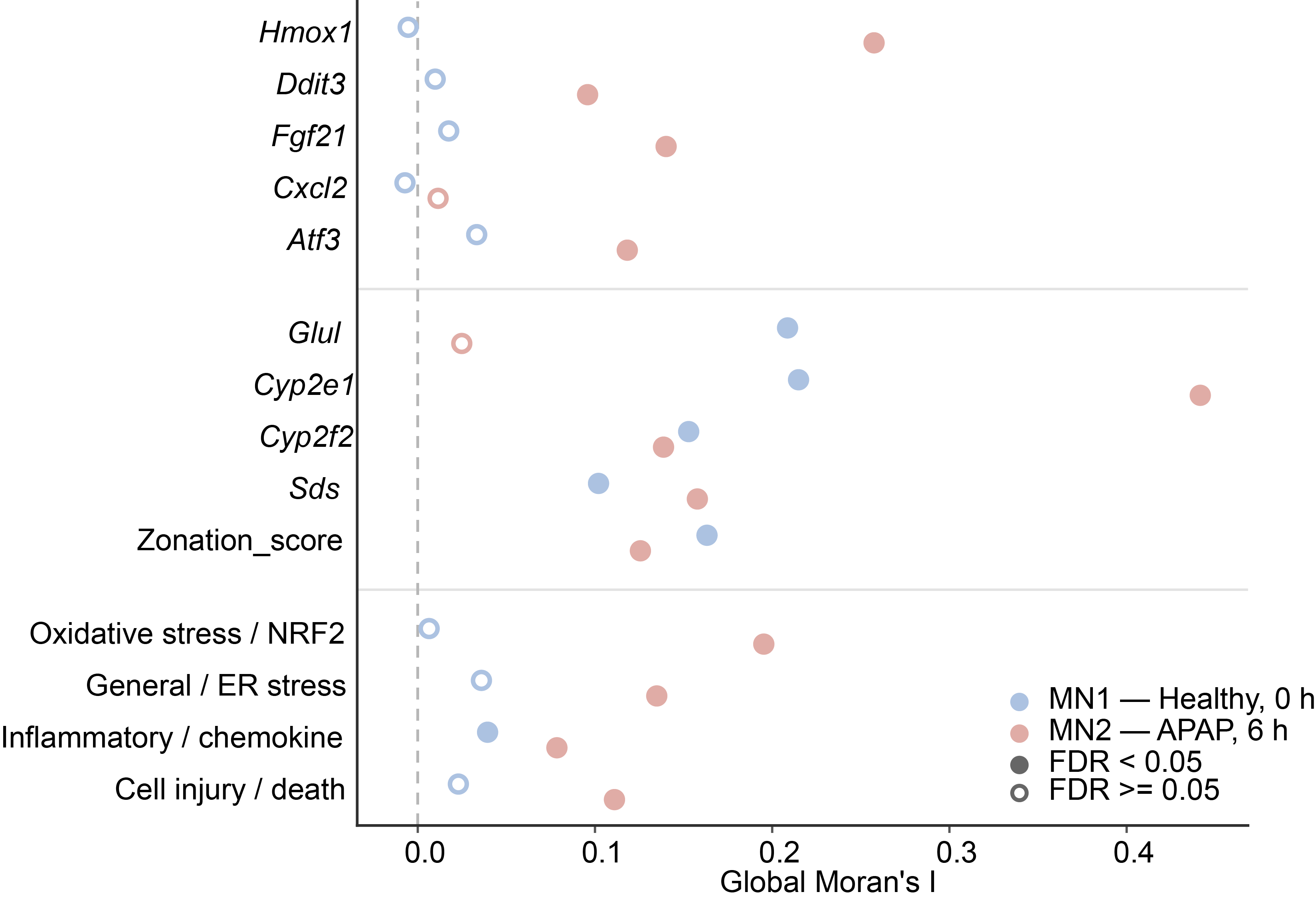

**Supplementary Fig. 9 Global spatial autocorrelation of representative APAP-responsive genes, zonation-associated genes, and transcriptional modules.** Global Moran’s I values are shown for representative APAP/stress-related genes, liver zonation-associated genes and the zonation score, and four APAP injury-related transcriptional modules in MN1 and MN2. Filled circles indicate significant global spatial autocorrelation after multiple-testing correction (false discovery rate [FDR] < 0.05), whereas open circles indicate FDR ≥ 0.05.

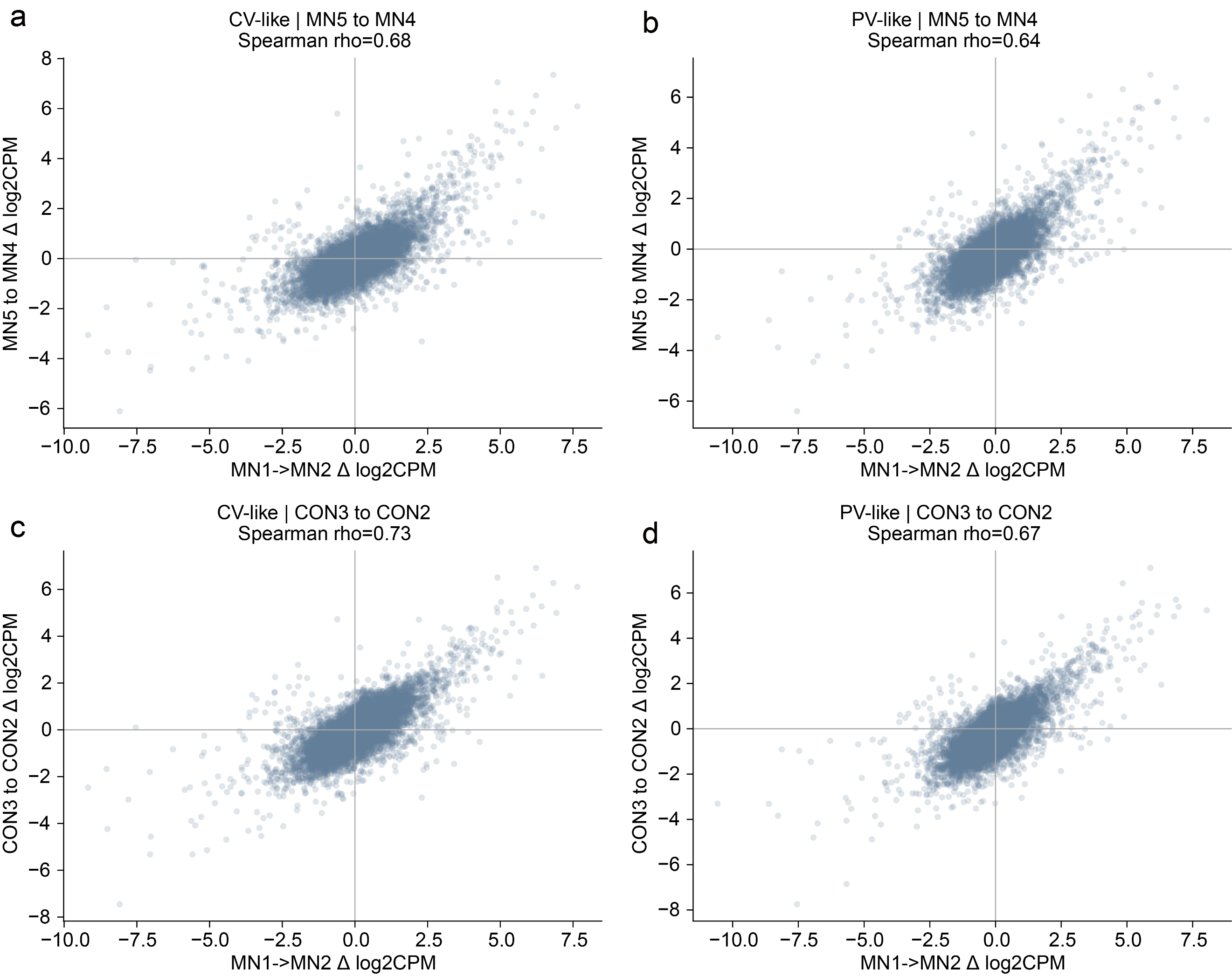

**Supplementary Fig. 10 Supportive concordance of zone-stratified transcriptional changes across liver samples.** (a, b) Gene-level MN1-to-MN2 changes compared with MN5-to-MN4 changes in CV-like (a) and PV-like (b) states. (c, d) Corresponding comparisons between MN1-to-MN2 and CON3-to-CON2 changes in CV-like (c) and PV-like (d) states. Spearman correlation coefficients are shown in each panel.

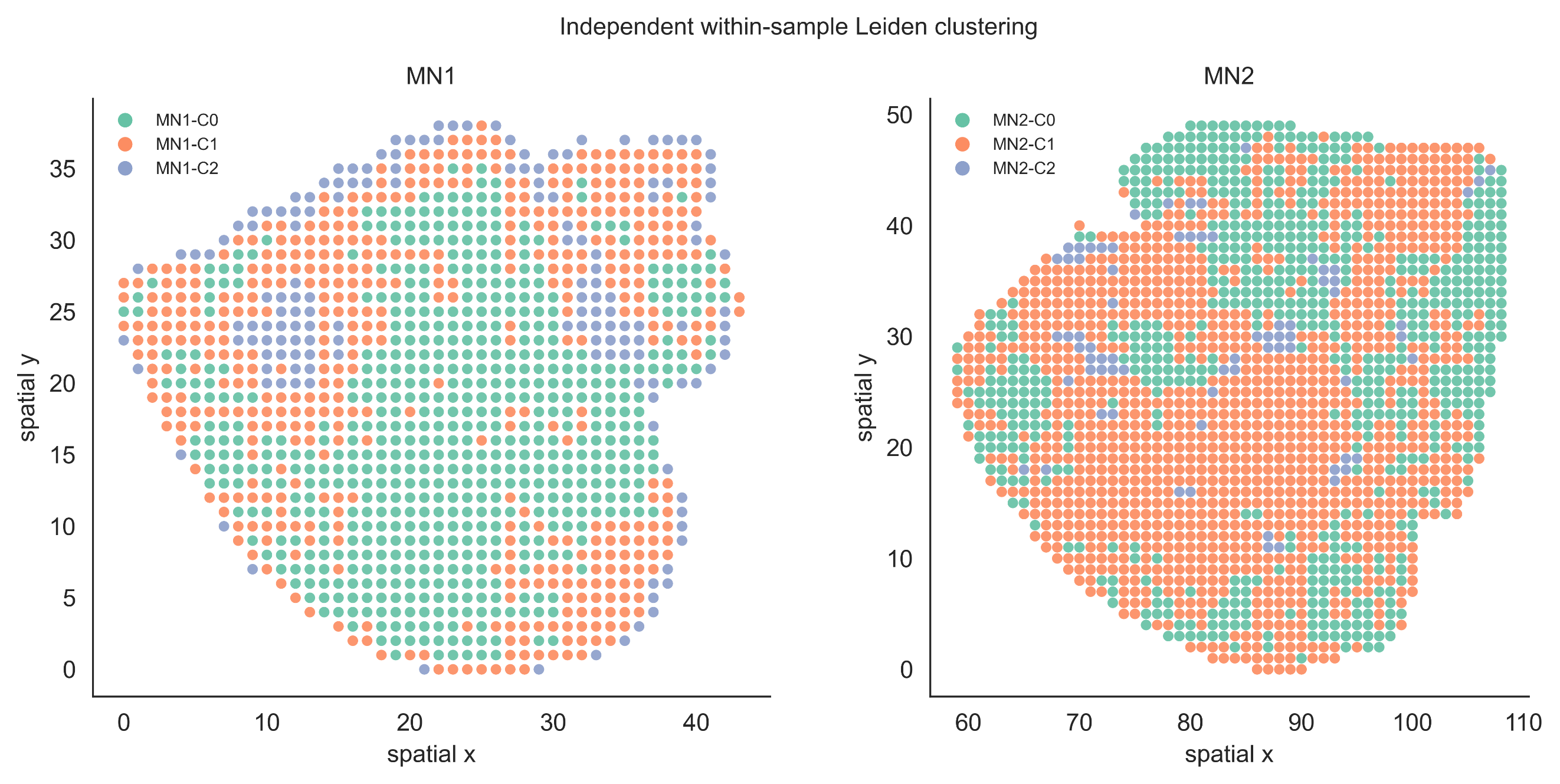

**Supplementary Fig. 11 Independent within-sample Leiden clustering of MN1 and MN2.** Leiden clustering was performed independently for MN1 and MN2, and the resulting cluster assignments were mapped back to their original spatial coordinates. Three transcriptional clusters were identified within each sample and showed partially interspersed spatial distributions.
